# Defective endosome-TGN retrograde transport promotes NLRP3 inflammasome activation

**DOI:** 10.1101/2021.09.14.460331

**Authors:** Zhirong Zhang, Li Ran, Rossella Venditti, Zengzhen Liu, Annette Schürmann, Juan S. Bonifacino, Maria Antonietta De Matteis, Romeo Ricci

## Abstract

Inflammasome complexes are pivotal in the innate immune response to pathogens and other danger signals^1–4^. The NLRP3 inflammasome is activated in response to a broad variety of cellular stressors. Most of the stimuli act in a potassium efflux-dependent manner but a primary and converging sensing mechanism by the NLRP3 receptor initiating inflammasome assembly remains ill-defined. Here we show that NLRP3 activators disrupt endosome-TGN retrograde transport (ETRT) and lead to localization of NLRP3 to endosomal vesicles. Genetic and pharmacologic perturbation of ETRT leads to accumulation of phosphoinositol-4-phosphate (PI4P) in endosomes to which NLRP3 is recruited. Disruption of ETRT potentiates NLRP3 inflammasome activation in murine and human macrophages *in vitro*. Mice with defects in ETRT in the myeloid compartment are more susceptible to LPS-induced sepsis showing enhanced mortality and IL-1β serum levels as compared to control animals. Our study thus uncovers that changes in endocytic trafficking mediate NLRP3-dependent inflammatory responses.

---

Inflammasomes are cytosolic multimeric protein complexes that play critical roles in eliciting innate immune responses against pathogens and damage-associated signals. The NLR family pyrin domain containing protein 3 (NLRP3) inflammasome is unique in that it is capable of detecting a remarkable variety of danger signals and is therefore broadly implicated in different inflammatory diseases^1–4^. NLRP3 inflammasome activation requires two principal steps, priming through Toll-like or cytokine receptor signaling resulting in robust expression of the inflammasome components and their assembly upon exposure with NLRP3 inflammasome activating factors. Depending on the nature of these activating stimuli, several mechanisms can lead to assembly of the NLRP3 inflammasome. Most of them trigger potassium efflux to activate the NLRP3 inflammasome^5,6^. Yet, a primary sensing mechanism of the NLRP3 inflammasome receptor remains largely elusive. We recently reported that NLRP3 localized to foci close to Golgi membranes^7^. In the attempt to further characterize NLRP3 localization, we used HeLa cells ectopically expressing EGFP-tagged NLRP3 devoid of endogenous expression of the inflammasome adaptor protein ASC that is required for inflammasome assembly. We observed that treatment with nigericin, a potassium efflux-dependent NLRP3 inflammasome activator, caused re-localization of endogenous TGN46 from the TGN to scattered vesicles, and association of EGFP-tagged NLRP3 to these vesicles (Extended Data Fig. 1). In contrast, the distribution of other TGN proteins such as the golgins GCC1/GCC88, GCC2/GCC185, Golgin 97 and Golgin 245/p230 was not affected by treatment with nigericin as well as with the imiquimod-derived compound CL097, a potassium efflux-independent NLRP3 activator (Extended Data Fig. 2a, 2b, 3a-c), excluding defects in overall TGN integrity. Unlike the golgins, TGN46 is a transmembrane protein that cycles between the plasma membrane and TGN. Upon endocytosis, TGN46 is delivered to endosomes and reaches the TGN through endosome-TGN retrograde transport^8^. Thus, impaired retrograde transport of TGN46 and consequently localization of NLRP3 to endosomes could explain co-localization of NLRP3 and TGN46.

To address whether NLRP3 inflammasome activation affected endosome-TGN retrograde transport (ETRT), we next examined internalization of Cy3-conjugated Shiga toxin subunit B (StxB) in HeLa cells in response to nigericin or CL097 treatment. Shiga toxins belong to a family of protein toxins that transit along the retrograde pathway^9,10^. We observed that in control-treated cells, the majority of StxB localized to a GCC1-positive TGN compartment after 20 and 40 min of endocytosis, in line with efficient endosomal retrograde transport (Fig. 1a, b). In contrast, in nigericin-treated (Fig. 1a, b) or CL097-treated cells (Extended Data Fig. 4a, b), StxB was mainly found on enlarged early endosome antigen 1 (EEA1)-positive vesicles. As expected, StxB-enriched EEA1-positive vesicles were neither positive for GCC1, GCC2 nor p230, confirming their endosomal origin (Fig. 1a and Extended Data Fig. 4a, c). Importantly, StxB co-localized with TGN46 upon nigericin treatment in line with the fact that TGN46 also transits along the endosome-TGN retrograde pathway (Extended Data Fig. 4d). These results indicated that treatment with nigericin or CL097 blocked endosome-TGN retrograde transport, leading to retention of StxB as well as TGN46 on endosomes. This is in line with previous studies showing that nigericin at least partially alleviates Shiga toxin related effects^11,12^. To corroborate this observation, we further assessed retrograde transport of the endogenous cation-independent mannose-6-phosphate receptor (CI-MPR) as previously described^13,14^. To this end, we examined the internalization of an antibody to the extracellular domain of CI-MPR^15^ in control-versus nigericin- or CL097-treated HeLa cells. The intracellular distribution of the CI-MPR antibody upon endocytosis was visualized by immunofluorescence staining. In control-treated cells, the CI-MPR antibody mainly localized to the TGN with only small amounts still at endosomes (Fig. 1b and Extended Data Fig. 5). In contrast, most of the CI-MPR antibody was detected on endosomes but not at the TGN in nigericin-as well as CL097-treated cells (Fig. 1b and Extended Data Fig. 5). These observations indicate that NLRP3 inflammasome activation impairs ETRT. The co-localization of TGN46 with NLRP3 may thus be indicative of recruitment of NLRP3 to endosomal vesicles. Indeed, we found NLRP3 on vesicles positive for the endosomal marker RAB5 (Fig. 2a). To characterize the dynamics of endosomal localization of NLRP3, we next used time-lapse spinning-disk microscope imaging in HeLa cells stably expressing mApple-tagged RAB5 and EGFP-tagged NLRP3. We observed that, upon addition of nigericin, NLRP3 rapidly formed small foci close to the plasma membrane that progressively became positive for RAB5, confirming the recruitment of NLRP3 to endocytic vesicles (Fig. 2b). These vesicles eventually develop into larger spheres at later time points of nigericin stimulation (Fig. 2b). Similar results were obtained in cells treated with CL097 (Extended Data Fig. 6). These results agree with a previous report that NLRP3-positive vesicles are positive for EEA1 and TGN46^16^. In this report, however, the authors concluded that EEA1 was missorted to Golgi-derived TGN46-positive vesicles. In contrast, our data support a model in which NLRP3-positive vesicles are of endosomal origin and that TGN46 is enriched on endosomes because of a defect in ETRT.

**Fig. 1.**
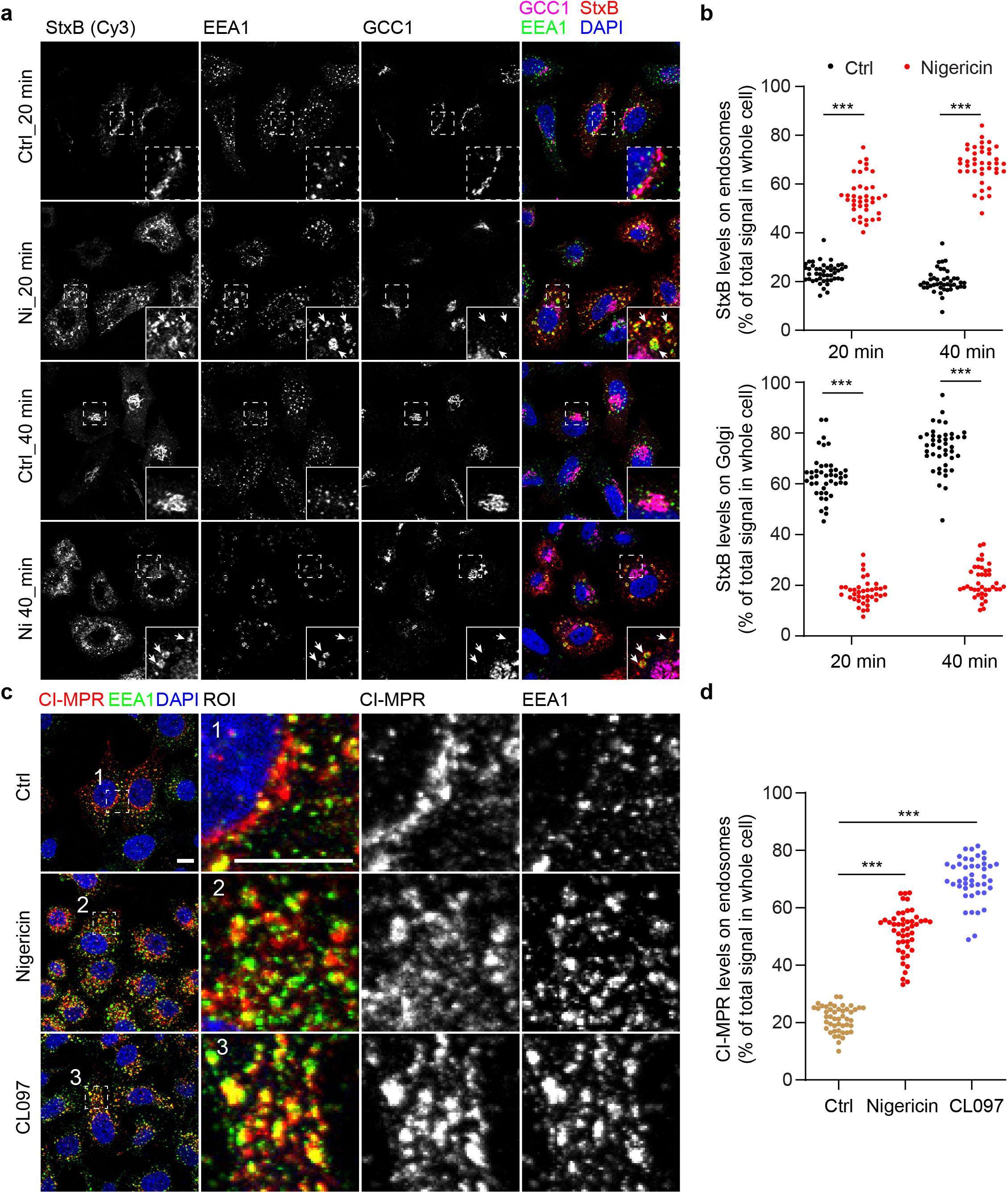
Endosome-TGN retrograde transport is inhibted in response to NLRP3 inflammasome activators. **a**, Internalization of shiga toxin subunit B (StxB) in HeLa cells. Cells were treated or not with 10 µM nigericin. Cy3-conjugated StxB was added into culture medium at 5 min after addition of nigericin. Cells were fixed at 20 min or 40 min after incubation with Cy3-conjugated StxB. Cells were stained with antibodies against GCC1 and EEA1. DAPI was used to stain the nucleus. Magnifications of areas are indicated with dashed squares and shown in the lower right corner of images. Scale bar: 10 µm. **b**, Quantification of StxB levels on endosomes and Golgi in experi-ments shown in Panel a. \*\*\**p*<0.0001 (around 45 cells per condition counted). **c**, Internalization of CI-MPR antibody in HeLa cells. Cells were treated or not with 10 µM nigericin or 45 µg/ml CL097. An antibody recognizing CI-MPR was added into the culture medium at 5 min after nigericin or at 20 min after CL097 treatment. Cells were fixed at 40 min after the incubation with the CI-MPR antibody and were stained with an antibody against EEA1. DAPI was used to stain the nucleus. Magnifications of areas are indicated with numbered dashed squares and shown in seperate images with corresponding numbers. Scale bar: 10 µm. **d**, Quantification of CI-MPR levels on endosomes in experiments shown in Panel c. \*\*\**p*<0.0001 (around 45 cells per condition counted). **a, c**, images are representative of at least three independent experiments.

**Fig. 2.**
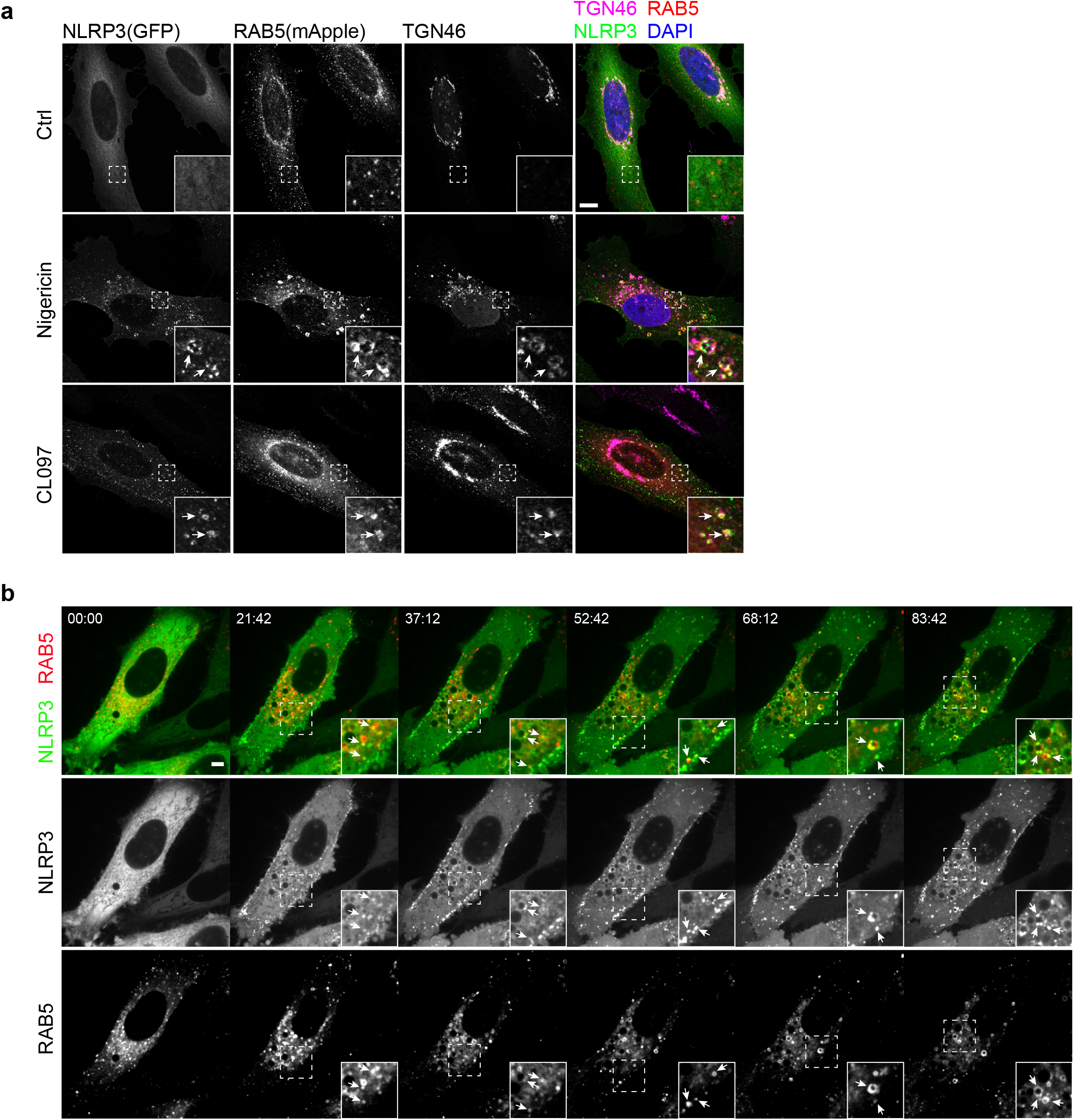
NLRP3 is recruited to vesicles of endosomal origin upon inflammasome activation. **a**, Confocal microscopy of HeLa cells stably expressing EGFP-tagged NLRP3 and mApple-tagged RAB5A. Cells were treated with vehicle (Ctrl), 10 µM Nigericin for 40 min or 45 µg/ml CL097 for 80 min. After fixation, cells were stained with an antibody against TGN46. DAPI was used to stain the nucleus. Arrows indicate triple-positive vesicles. Magnifications of areas are indicated with dashed squares and shown in the lower right corner of images. Scale bar: 10 µm. **b**, Spinning-disk confocal time-lapse live-video microscopy of HeLa cells stably expressing EGFP-tagged NLRP3 and mApple-tagged RAB5A. Live images were captured at every 186 seconds after addtion of 10 µM Nigericin into culture medium. Representative video stills at indicated time points are shown. Arrows indicate NLRP3 and Rab5 double-positive vesicles. Magnifications of areas are indicated with dashed squares and shown in the lower right corner of images. Scale bar: 10 µm. Images shown are representative of at least three independent experiments.

To further examine the relationship of ETRT and NLRP3 inflammasome activation, we disrupted ETRT through genetic means and assessed the effects of this manipulation on NLRP3 inflammasome activation. Both ADP ribosylation factor related protein 1 (ARFRP1) and SYS1 are critical regulators of TGN tethering factors, including long coiled-coil tethers of the golgin family and the heterotetrameric complex GARP, which mediates fusion of endosome-derived transport carriers with the TGN membranes ^17^. Therefore, we deleted the *ARFRP1* and *SYS1* genes in THP-1 cells, a human acute leukemia monocytic cell line widely used for assessing NLRP3 inflammasome activation, using CRISPR/Cas9-mediated gene editing. Gene disruption was validated by immunoblotting or Sanger sequencing (Extended Data Fig. 7a-c). In line with a previous report^17^, deletion of *SYS1* reduced protein levels of ARFRP1 (Extended Data Fig. 7c). Deletion of *ARFRP1* or *SYS1* did not affect expression of the inflammasome components, NLRP3, ASC, Caspase 1 and NEK7 (Extended Data Fig. 7c). However, priming through stimulation with lipopolysaccharide (LPS) (a TLR4 ligand) or Pam3CSK4 (a TLR2/TLR1 ligand) was sufficient to induce lytic cell death of *ARFRP1* knockout (KO) and *SYS1* KO THP-1 cells, as evidenced by the uptake of the live cell-impermeant nucleic acid dye Sytox Green (Fig. 3a), accompanied by the cleavage and secretion of Caspase 1 and IL-1β (Extended Data Fig. 8a, b). In contrast, LPS or Pam3CSK4 alone did not evoke such a response in parental wild type (WT) cells (Fig. 3a and Extended Data Fig. 8a, b). LPS-induced lytic cell death, cleavage and secretion of Caspase 1 and IL-1β in KO cells were dependent on NLRP3, as these effects were abolished in the presence of the NLRP3 inhibitor MCC950 (Fig. 3b and Extended Data Fig. 8a, b) or by deletion of NLRP3 (Extended Data Fig. 8c). LPS treatment induced the assembly of mature inflammasome in these KO cells as evidenced by the formation of ASC specks (Extended Data Fig. 8d, e). LPS-induced NLRP3 inflammasome activation in KO cells was independent of potassium efflux, as high extracellular potassium levels did not affect Caspase 1 cleavage induced by LPS (Fig. 3b). Furthermore, re-expression of WT ARFRP1 or constitutively active ARFRP1 Q79L, but not inactive ARFRP1 T31N or FLAG-tagged SYS1 abrogated LPS-induced NLRP3 inflammasome activation in *ARFRP1* KO THP-1 cells (Fig. 3c, d), whereas only re-expression of FLAG-tagged SYS1, but not ARFRP1 WT, Q79L or T31N, prevented LPS-induced NLRP3 inflammasome activation in *SYS1* KO THP-1 cells (Fig. 3c, d). These results are in line with previous studies showing that SYS1 and ARFRP1 are both critical components of TGN tethering^17,18^. From these experiments, we concluded that perturbation of the TGN tethering machinery was sufficient to activate the NLRP3 inflammasome in response to priming in THP-1 cells in a potassium efflux-independent manner.

**Fig. 3.**
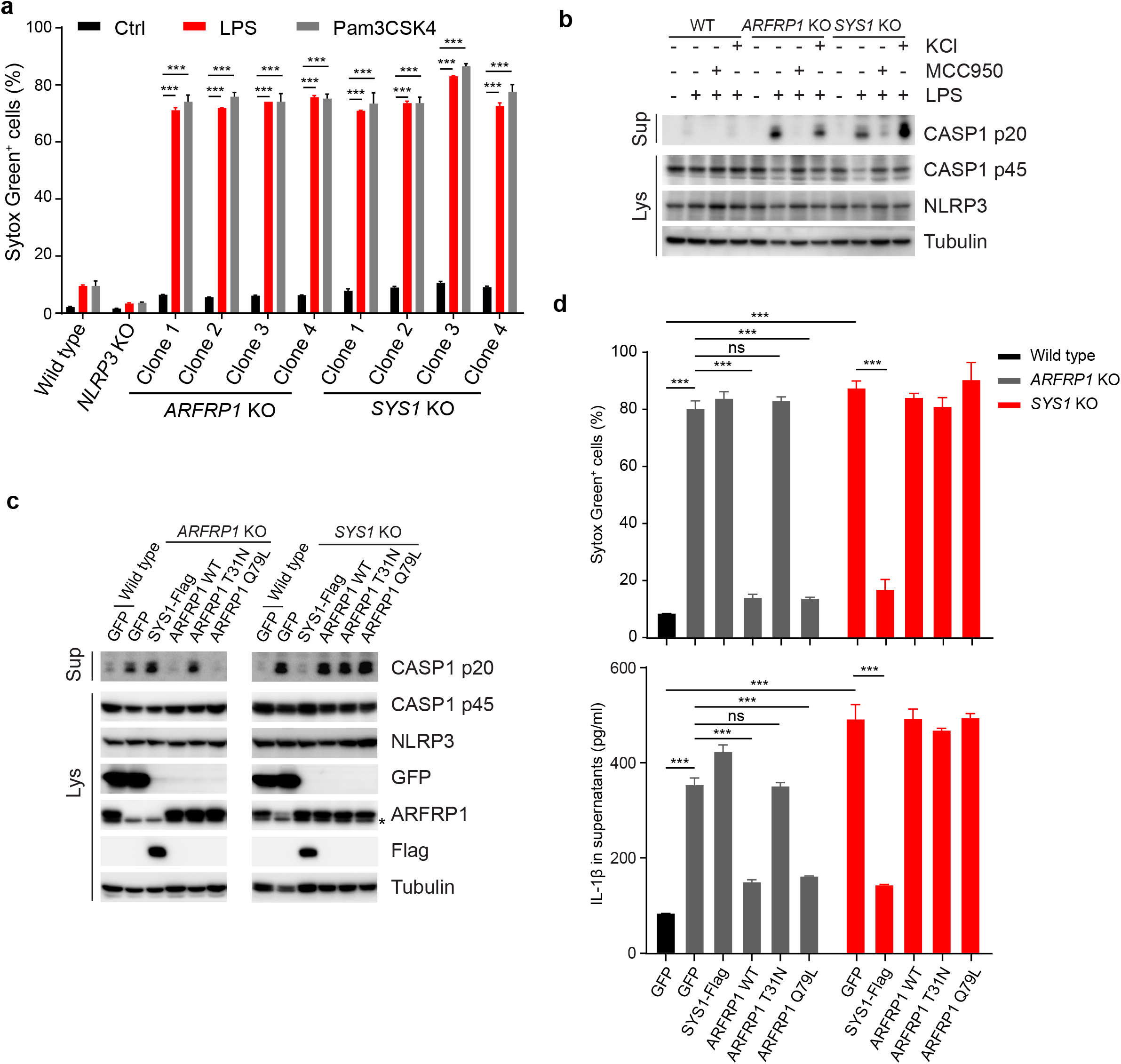
Genetic disruption of endosome-TGN retrograde transport induces activation of the NLRP3 inflammasome in response to priming only. **a**, Cellular uptake of Sytox Green in PMA-differentiated THP-1 cells treated with LPS or Pam3CSK4. Wild type, *NLRP3* KO, *ARFRP1* KO and *SYS1* KO THP-1 cells were treated or not with 1 µg/ml LPS or Pam3CSK4 for 2 hours. Cells were stained with Sytox Green and subjected to FACS analysis. \*\*\**p* < 0.001. **b**, Immunoblotting of culture supernatants and cell lysates from THP-1 cells. Wild type, *ARFRP1* KO and *SYS1* KO THP-1 cells were treated or not with 1 µg/ml LPS in absence or presence of the NLRP3 inhibitor, MCC950 (10 µM) or KCl (20 mM) for 2 hours. Antibodies recognizing both p45 and p20 fragments of Caspase 1 (CASP1), IL1β and NLRP3 were used. An antibody against Tubulin was used as a loading control. **c**, Immunoblotting of culture supernatants and cell lysates from THP-1 cells. Wild type, *ARFPR1* KO (left) and *SYS1* KO (right) THP-1 cells expressing GFP, Flag-tagged SYS1, wild type ARFRP1 (ARFRP1 WT), T31N mutant ARFRP1 (ARFRP1 T31N) or Q79L mutant ARFRP1 (ARFRP1 Q79L) were treated with 1 µg/ml LPS for 2 hours. Antibodies against CASP1, NLRP3, GFP, ARFRP and Flag were used. An antibody against Tubulin was used as a loading control. **d**, Uptake of Sytox Green by FACS (upper panel) and ELISA assay of IL1β in culture supernatants (lower panel) of THP-1 cells. Wild type, *ARFPR1* KO (left) and *SYS1* KO (right) THP-1 cells expressing GFP, Flag-tagged SYS1, wild type ARFRP1 (ARFRP1 WT), T31N mutant ARFRP1 (ARFRP1 T31N) or Q79L mutant ARFRP1 (ARFRP1 Q79L) were treated with 1 µg/ml LPS for 2 hours. \*\*\**p* < 0.001; not significant (ns). Data shown are representative of at least three independent experiments.

Retro-2 is a chemical compound that selectively blocks retrograde trafficking of ricin and Shiga-like toxins at the early endosome-TGN interface^19,20^. Therefore, we next assessed NLRP3 inflammasome activation in response to NLRP3 activators in the presence or absence of Retro-2. Retro-2 treatment was not sufficient to induce NLRP3 inflammasome activity in THP-1 cells in suspension upon priming only (Extended Data Fig. 9)^19^. However, treatment with Retro-2 significantly enhanced NLRP3 inflammasome activity, as evidenced by increased Caspase 1 activation, IL-1β cleavage and secretion in PMA-differentiated THP-1 cells stimulated with nigericin or CL097 (Extended Data Fig. 9a, b). Likewise, Retro-2-treated mouse primary bone marrow-derived macrophages (BMDMs) showed a significantly increased IL-1β release in response to ATP, nigericin, CL097 and imiquimod (R837) (Extended Data Fig. 9c).

To further test whether this mechanism is conserved among species, we next generated immortalized mouse bone marrow-derived macrophages (iBMDMs) lacking *Arfrp1* or *Sys1* and tested NLRP3 inflammasome activation. As observed in THP-1 KO cells, both *Arfrp1* and *Sys1* KO iBMDMs showed a significant IL-1β release in response to LPS priming in the absence of NLRP3 activators, while WT cells did not (Extended Data Fig. 10). However, KO iBMDMs did not undergo lytic cell death in response to LPS priming as observed in THP-1 cells. Yet, KO iBMDMs revealed dramatically enhanced NLRP3 inflammasome activation when treated with nigericin or CL097 as compared to WT cells (Extended Data Fig. 10).

Together with the above results, these data suggest that inhibition of ETRT as well as recruitment of NLRP3 to endosomes in response to potassium-dependent and -independent NLRP3 inflammasome activators mediate NLRP3 inflammasome activation.

NLRP3 was shown to bind to phosphatidylinositol 4-phosphate (PI4P)^16^. We thus next wondered whether PI4P levels in the endosomal compartment increase in response to inflammasome activators. Indeed, using an antibody against PI4P, we observed that PI4P levels on endosomes dramatically increased (Extended Data Fig. 11a-d). Furthermore, nigericin-induced NLRP3 foci strongly co-localized with PI4P (Extended Data Fig. 11e). Phosphatidylinositol 4-kinase IIIβ (PI4KIIIβ) is a main enzyme generating PI4P in the TGN^21,22^. Indeed, deletion of PI4KIIIβ efficiently depleted PI4P in TGN membranes (Extended Data Fig. 12). However, this did not affect enrichment of both PI4P and NLRP3 on endosomes upon nigericin or CL097 stimulation (Extended Data Fig. 12b), further supporting that the source of NLRP3-positive vesicles and PI4P is not the TGN. Finally, perturbed TGN tethering activity in *ARFRP1* KO and *SYS1* KO HeLa cells led to accumulation of PI4P on endosomes (Extended Data Fig. 13a, b) analogous to what we observed in cells in response to NLRP3 inflammasome activators. Similar results were obtained in *ARFRP1* KO and *SYS1* KO THP-1 cells (Extended Data Fig. 13c). Altogether, these results indicate that NLRP3 activators disrupt ETRT, leading to accumulation of PI4P on endosomes and endosomal NLRP3 recruitment to activate the NLRP3 inflammasome.

To assess the *in vivo* relevance of our findings, we next generated mice lacking *Arfrp1* in the myeloid lineage by crossing *Arfrp1* floxed mice to *LysM-cre* mice. Efficient deletion of Arfrp1 in BMDMs was confirmed by Western blotting (Fig. 4a). *Arfrp1* KO BMDMs displayed significantly enhanced Caspase1 activation, IL-1β cleavage as well as IL-1β secretion in response to ATP, nigericin, CL097 or R837 treatment compared to control cells (Fig. 4a, b). While the secretion of TNFα in these cells is comparable to that in WT BMDMs (Fig. 4b). NLRP3 inflammasome activation and IL-1β release is crucial in low dose LPS-induced endotoxemia^23^. Thus, we next performed intraperitoneal injections using sublethal doses of LPS. Strikingly, myeloid-specific *Arfrp1* KO mice were significantly more susceptible to an LPS challenge as compared to control mice as most of the myeloid-specific *Arfrp1* KO mice died, while most control mice survived such doses of LPS (Fig. 4c). In line with enhanced NLRP3 inflammasome activation, serum IL-1β levels were significantly enhanced in myeloid-specific *Arfrp1* KO mice as compared to control mice (Fig. 4d); while serum TNFα levels were comparable between control mice and myeloid-specific *Arfrp1* KO mice (Fig. 4d). These results suggest that defective ETRT drives NLRP3 inflammasome activation *in vivo* in mice.

**Fig. 4.**
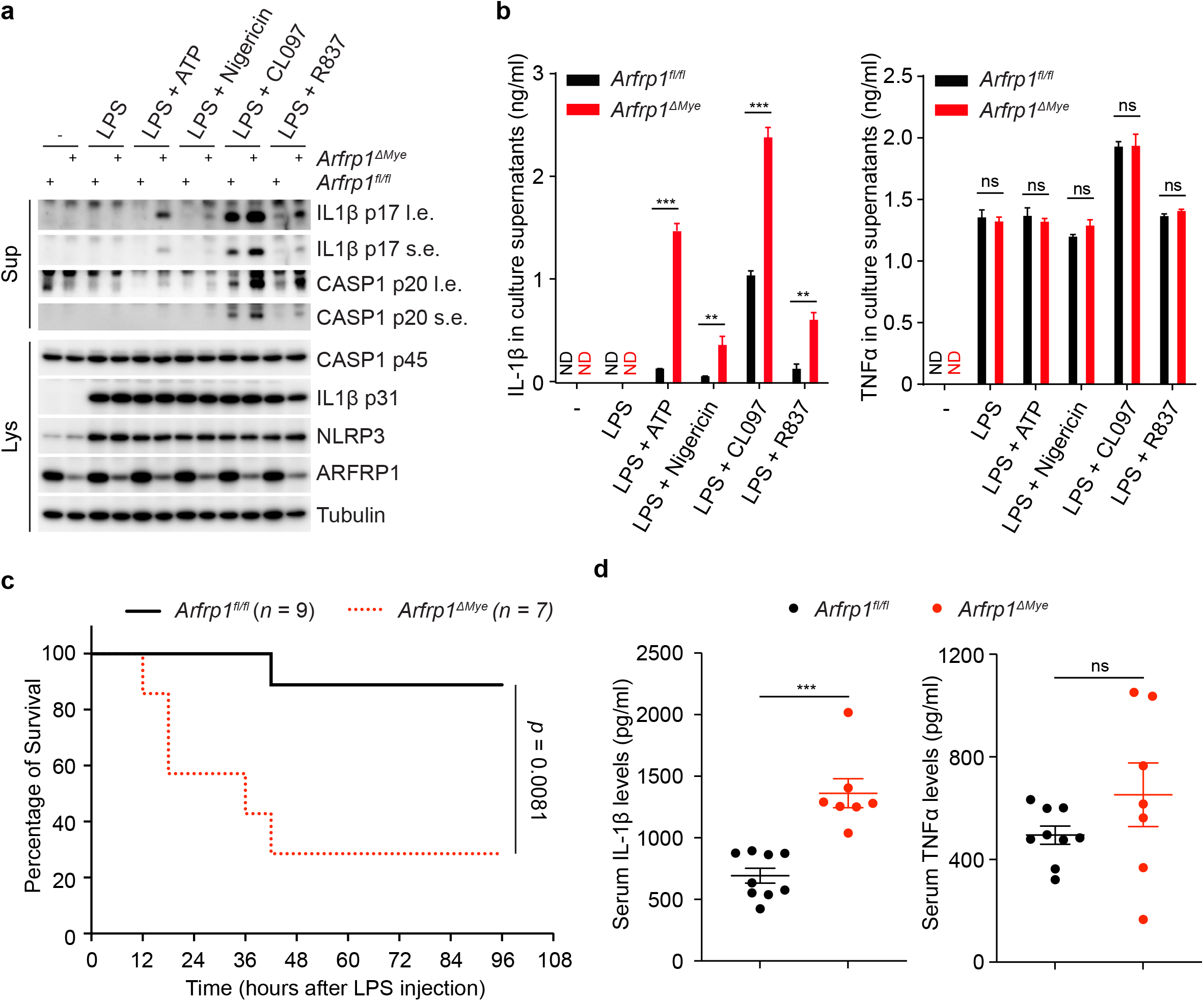
Genetic disruption of endosome-TGN retrograde transport promotes NLRP3 inflammasome activation *in vivo* in mice. **a**, Immunoblotting of culture supernatants and cell lysates from primary BMDMs isolated from *Arfrp1*^*fl/fl*^ or *Arfrp1*^*fl/-fl*^;*LysM-Cre* (*Arfrp1*^*ΔMye*^) mice. Isolated cells were primed with 1 µg/ml LPS for 4 hours, followed by treatment with 2 mM ATP, 5 µM Nigericin, 50 µM CL097 or 50 µM R837 for 40 min as indicated. Antibodies recognizing both p45 and p20 fragments of Caspase 1 (CASP1), IL1β, NLRP3 and ARFRP1 were used. An antibody against Tubulin was used as a loading control. Short exposures (s.e.) and long exposures (l.e.) of IL1β and CASP1 fragments are shown. **b**, ELISA assay of IL1β and TNFα in culture supernatants of isolated BMDMs from mice as indicated. not detected (ND); \*\**p* < 0.01 and \*\*\**p* < 0.001. **c**, Survival curves of *Arfrp1*^*fl/fl*^ and *Arfrp1*^*fl/fl*^;*LysM-Cre* (*Arfrp1*^*ΔMye*^) mice injected with LPS (15 mg/kg). The survival of mice after LPS injection was monitored every 6 hours. *p* = 0.0081. **d**, ELISA assay of IL1β (left panel) and TNFα (right panel) levels in serum collected from *Arfrp1*^*fl/fl*^ or *Arfrp1*^*fl/fl*^;*LysM-Cre* (*Arfrp1*^*ΔMye*^) mice at 3 hours after peritoneal injections of LPS (15 mg/kg). \*\*\**p* < 0.001; not significant (ns). **a, b**, Data shown are representative of at least three independent experiments.

The NLRP3 inflammasome receptor is rather exceptional as it is capable of sensing very broadly cellular stress. We demonstrate here that a broad array of NLRP3 inflammasome activators converge into defective ETRT. ETRT might be hampered at different levels: at the endosomal level, possibly by impairing the sorting of retrograde cargo into retrogradely directed tubules or at the level of retrograde carriers that detach from endosomes but that cannot fuse with the TGN. Importantly, perturbation of ETRT on its own was sufficient to render cells highly susceptible to NLRP3 inflammasome activation *in vitro* as well as *in vivo*. Furthermore, blocking of ETRT results in accumulation of PI4P in endosomes, a distinct feature that we have also observed upon stimulation with NLRP3 inflammasome activators whether they are dependent on potassium efflux or not. Even though NLRP3 was shown to bind PI4P, it is not excluded that other factors are necessary for NLRP3 recruitment to endosomes. One potential explanation for endosomal accumulation of PI4P is concomitant disintegration of endosome-endoplasmic reticulum (ER) contacts, that has been shown to increase PI4P in endosomes as it is no longer transferred to the ER, where it should be eliminated by the phosphoinositide phosphatase Sac1^24^. In conclusion, our study provides evidence for NLRP3 to primarily sense endosomal stress occurring at least partially through defective ETRT.

## Acknowledgments

We acknowledge Dr. Anne Spang at the Biozentrum University of Basel, Switzerland, for providing HeLa cells stably expressing mApple-tagged RAB5. We thank Dr. Fabien Alpy, Dr. Catherine-Laure Tomasetto and Dr. Izabela Sumara for helpful suggestions and discussions. We also thank the scientific platforms at the IGBMC including the cell cytometry facility, the cell culture facility, the imaging facility and the animal facility for their help during the whole project.

## Author contributions

Conceptualization, Z.Z., R.R., M.A.D.M and J.S.B; Methodology, Z.Z., L.R., R.V. and Z.L.; Investigation, Z.Z., L.R., R.V., and Z.L.; Writing – Original Draft, Z.Z.; Writing – Review & Editing, R.R., Z.Z., J.S.B., A.S. and M.A.D.M.; Funding Acquisition, R.R.; Resources, Z.Z., E.E., J.S.B and A.S.; Supervision, R.R. and Z.Z.

## Data and materials availability

All data is available in the main text or the supplementary materials.

## Supplementary Materials

Extended Data Fig. 1-13

Methods

**Extended Data Fig.1.**
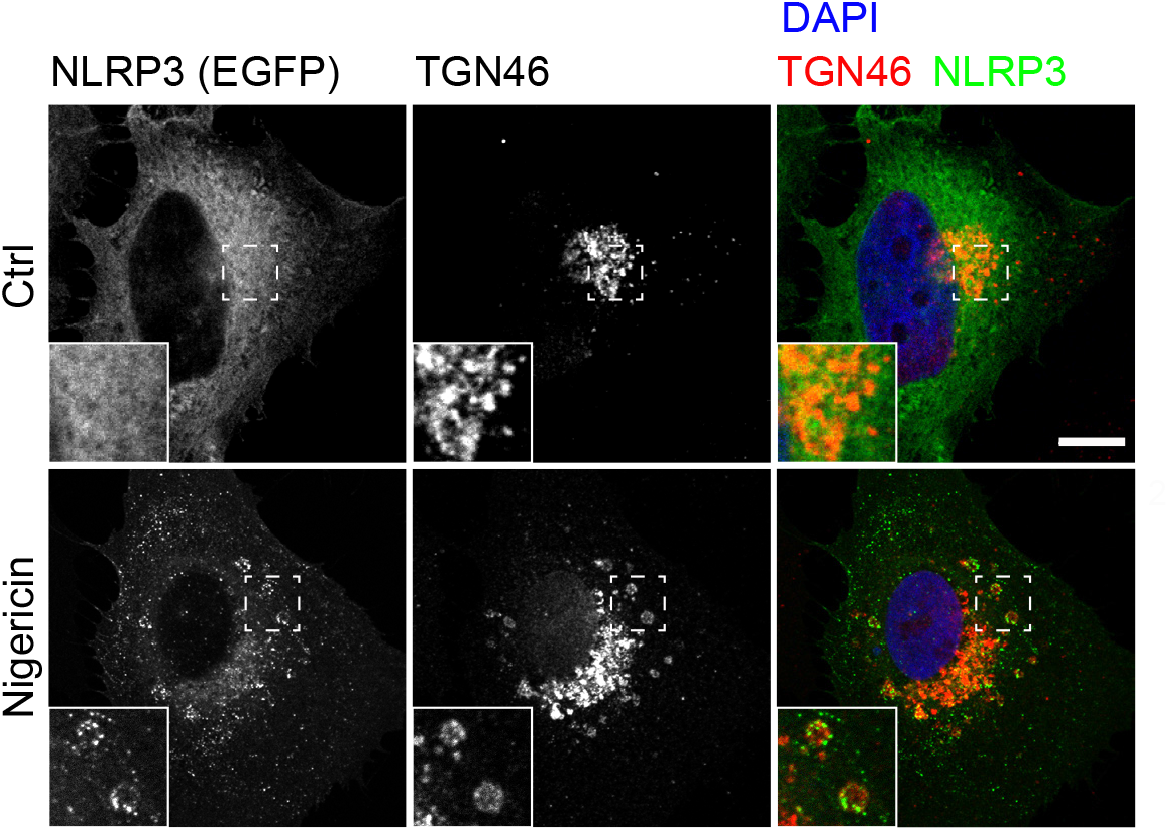
Nigericin treatment induces co-localization of NLRP3 with TGN46. Confocal microscopy of HeLa cells stably-expressing EGFP-tagged NLRP3 treated with vehicle or 10 µM Nigericin for 40 min. After fixation, cells were stained with an antibody against TGN46. DAPI was used to stain the nucleus. Magnifications of areas are indicated with dashed squares and shown in the lower left corner of images. Scale bar: 10 µm. Images shown are representative of three independent experiments.

**Extended Data Fig.2.**
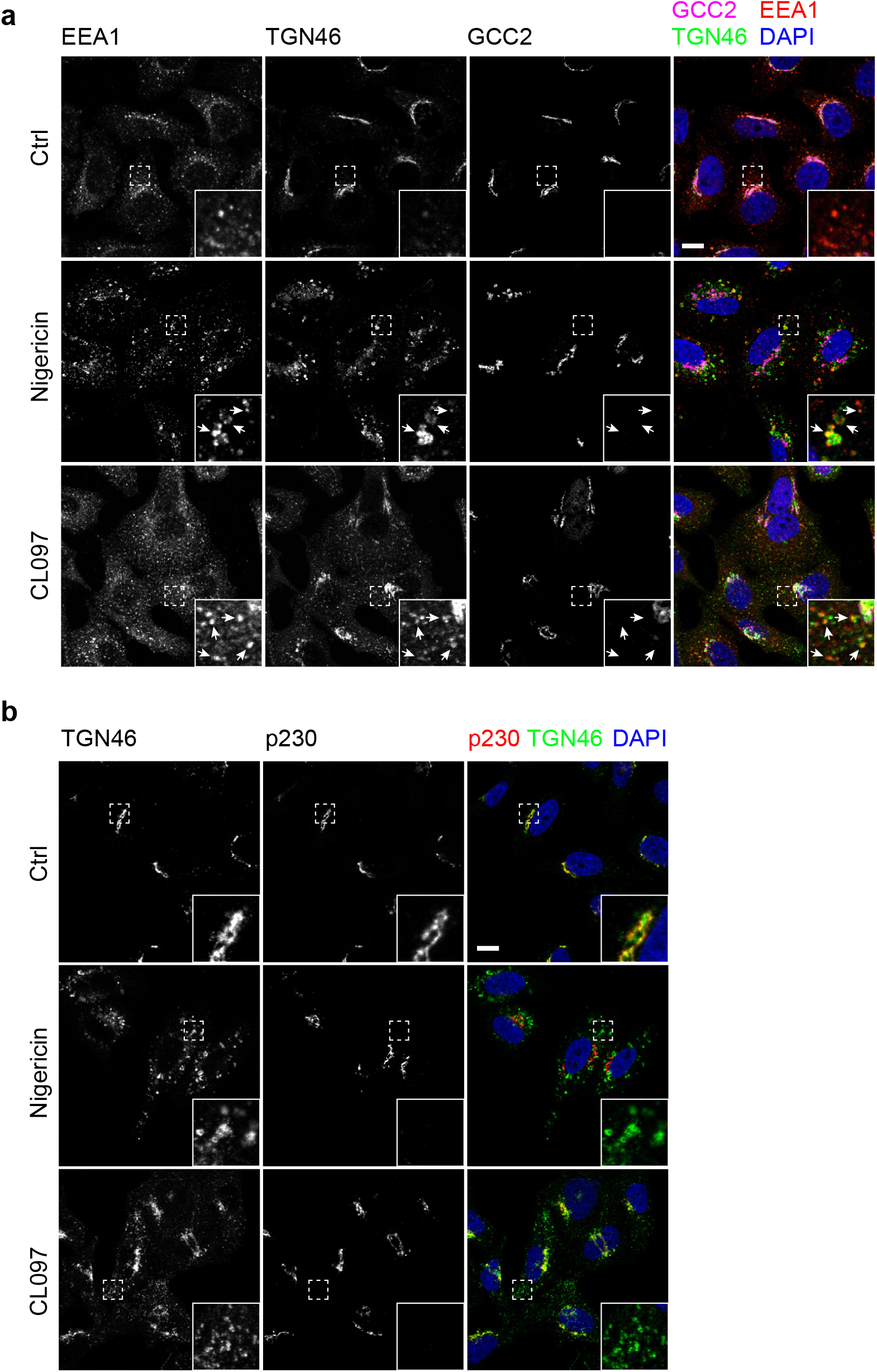
Both Nigericin and CL097 treatment redistribute exclusively TGN46 but not other Trans-Golgi network (TGN) proteins. **a**, Confocal microscopy of HeLa cells treated with vehicle, 10 µM Nigericin for 40 min or 45 µg/ml CL097 for 80 min. After fixation, cells were stained with antibodies against EEA1, TGN46 and GCC2. DAPI was used to stain the nucleus. Magnifications of areas are indicated with dashed squares and shown in the lower right corner of images. Scale bar: 10 µm. **b**, Confocal microscopy of HeLa cells treated with vehicle, 10 µM Nigericin for 40 min or 45 µg/ml CL097 for 80 min. After fixation, cells were stained with antibodies against TGN46 and p230. DAPI was used to stain the nucleus. Magnifications of areas are indicated with dashed squares and shown in the lower right corner of images. Scale bar: 10 µm. Images shown are representative of at least three independent experiments.

**Extended Data Fig.3.**
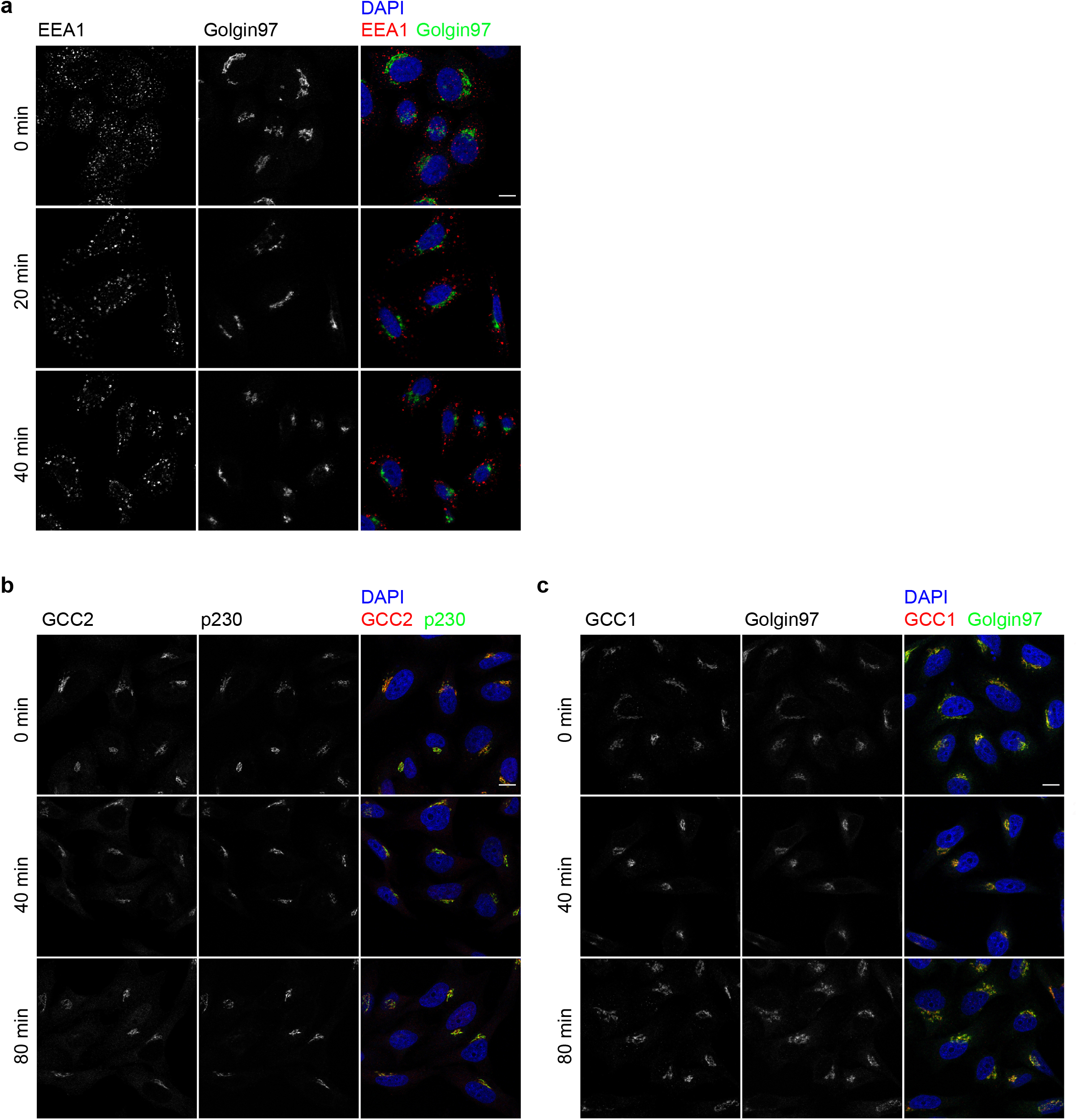
Neither Nigericin nor CL097 treatment induces overt Trans-Golgi network (TGN) dispersal. **a**, Confocal microscopy of HeLa cells treated with 10 µM Nigericin for 0 min, 20 min and 40 min. After fixation, cells were stained with antibodies against EEA1 and Golgin97. DAPI was used to stain the nucleus. Scale bar: 10 µm. **b**, Confocal microscopy of HeLa cells treated with 45 µg/ml CL097 for 0 min, 40 min and 80 min. After fixation, cells were stained with antibodies against GCC2 and p230. DAPI was used to stain the nuclear. **c**, Confocal microscopy of HeLa cells treated with 45 µg/ml CL097 for 0 min, 40 min and 80 min. After fixation, cells were stained with antibodies against GCC1 and Golgin97. DAPI was used to stain the nuclear. Scale bar: 10 µm. Images shown are representative of at least three independent experiments.

**Extended Data Fig.4.**
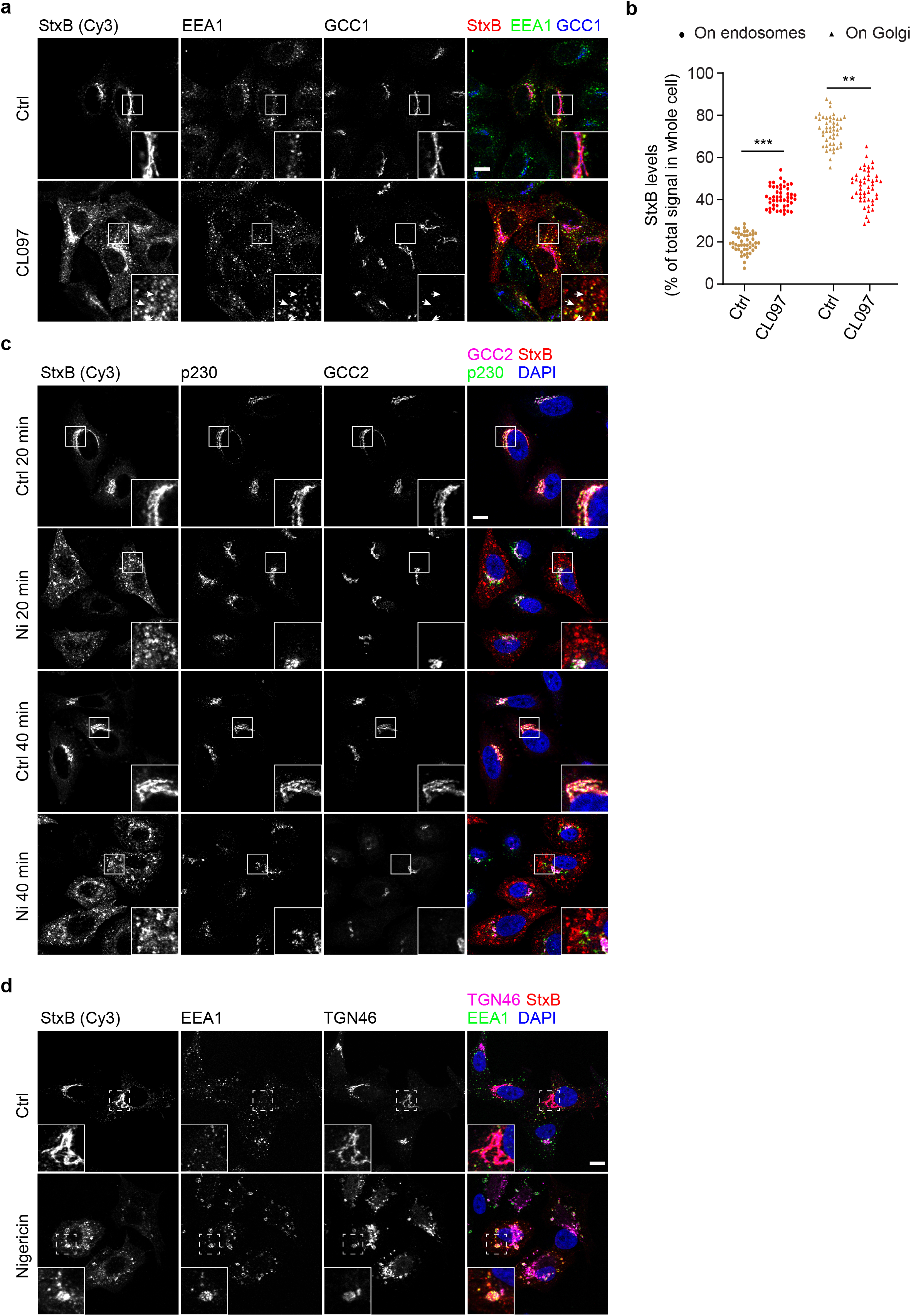
Endosome-TGN retrograde transport is inhibted in response to NLRP3 inflammasome activators. **a**, Internalization of StxB in HeLa cells. Cells were treated or not with 45 µg/ml CL097. Cy3-conjugated StxB was added into culture medium at 20 min after CL097 addition. Cells were fixed at 40 min after incubation with Cy3-conjugated StxB and stained with antibodies against EEA1 and GCC1. DAPI was used to stain the nucleus. Magnifications of areas are indicated with dashed squares and shown in the lower right corner of images. Scale bar: 10 µm. **b**, Quantification of StxB levels on endosomes and Golgi in experiments shown in panel **a**. ** *p* < 0.01; *** *p* < 0.001. **c**, Internalization of StxB in HeLa cells. Cells were treated or not with 10 µM Nigericin. Cy3-conjugated StxB was added into culture medium at 5 min after Nigericin addition. Cells were fixed at 20 min or 40 min after the incubation of Cy3-conjugated StxB and stained with antibodies against p230 and GCC2. DAPI was used to stain the nucleus. Magnifications of areas are indicated with dashed squares and shown in the lower right corner of images. Scale bar: 10 µm. **d**, Internalization of StxB in HeLa cells. Cells were treated or not with 10 µM Nigericin. Cy3-conjugated StxB was added into culture medium at 5 min after Nigericin addition. Cells were fixed at 40 min after the incubation with Cy3-conjugated StxB and stained with antibodies against EEA1 and TGN46. Magnifications of areas are indicated with dashed squares and shown in the lower left corner of images. DAPI was used to stain the nucleus. Scale bar: 10 µm. **a**,**c**,**d**, Images shown are representative of at least three independent experiments.

**Extended Data Fig.5.**
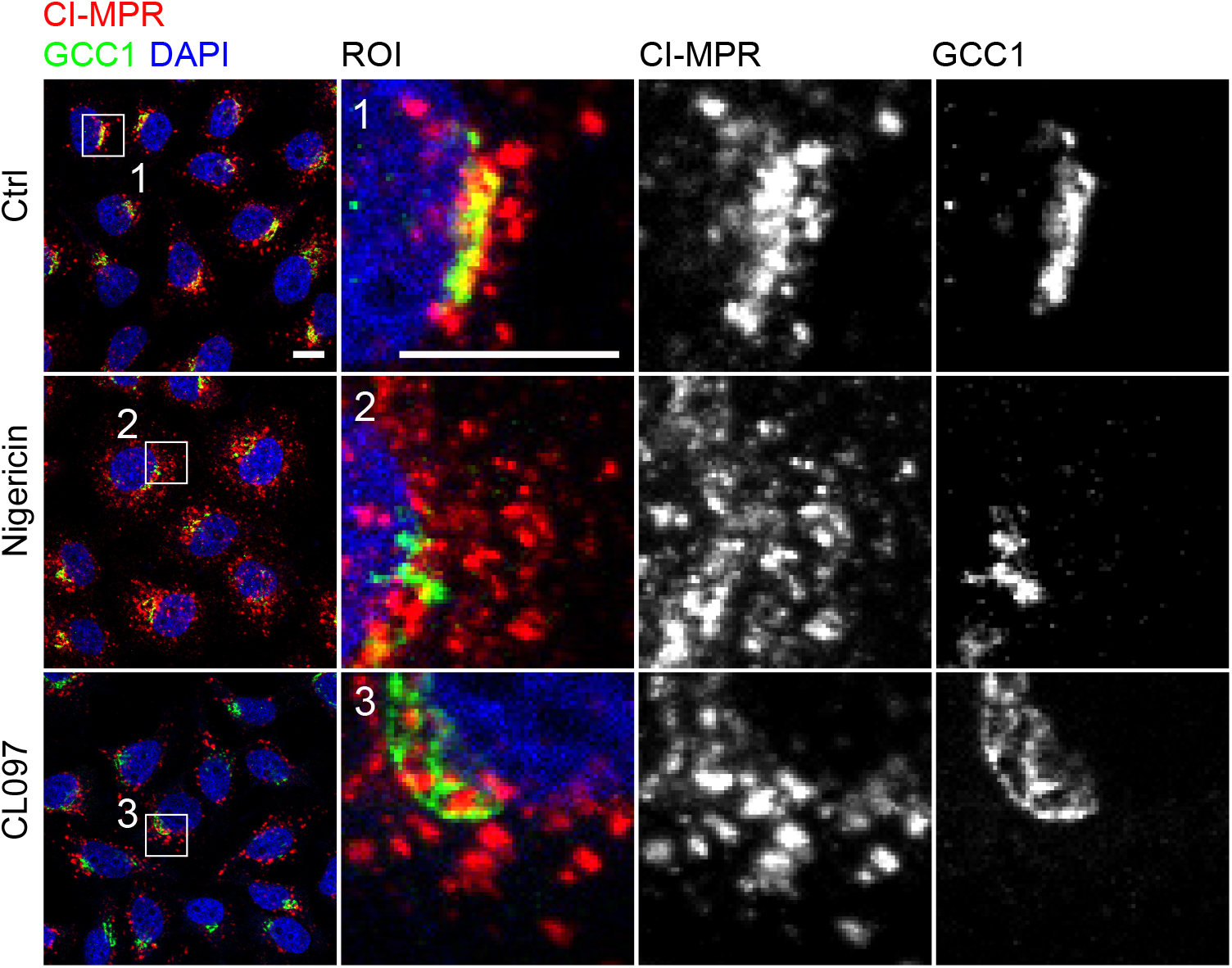
Endosome-TGN retrograde transport is inhibited in response to NLRP3 inflammasome activators. Internalization of CI-MPR antibody in HeLa cells. Cells were treated or not with 10 µM nigericin or 45 µg/ml CL097. An antibody recognizing CI-MPR was added into culture medium at 5 min after nigericin addition or at 20 min after CL097 addition. Cells were fixed at 40 min after the incubation with the CI-MPR antibody and were stained with antibodies against GCC1. DAPI was used to stain the nucleus. Magnifications of areas are indicated with numbered dashed squares and shown in seperate images with corresponding numbers. Scale bar: 10 µm. Images shown are representative of three independent experiments.

**Extended Data Fig.6.**
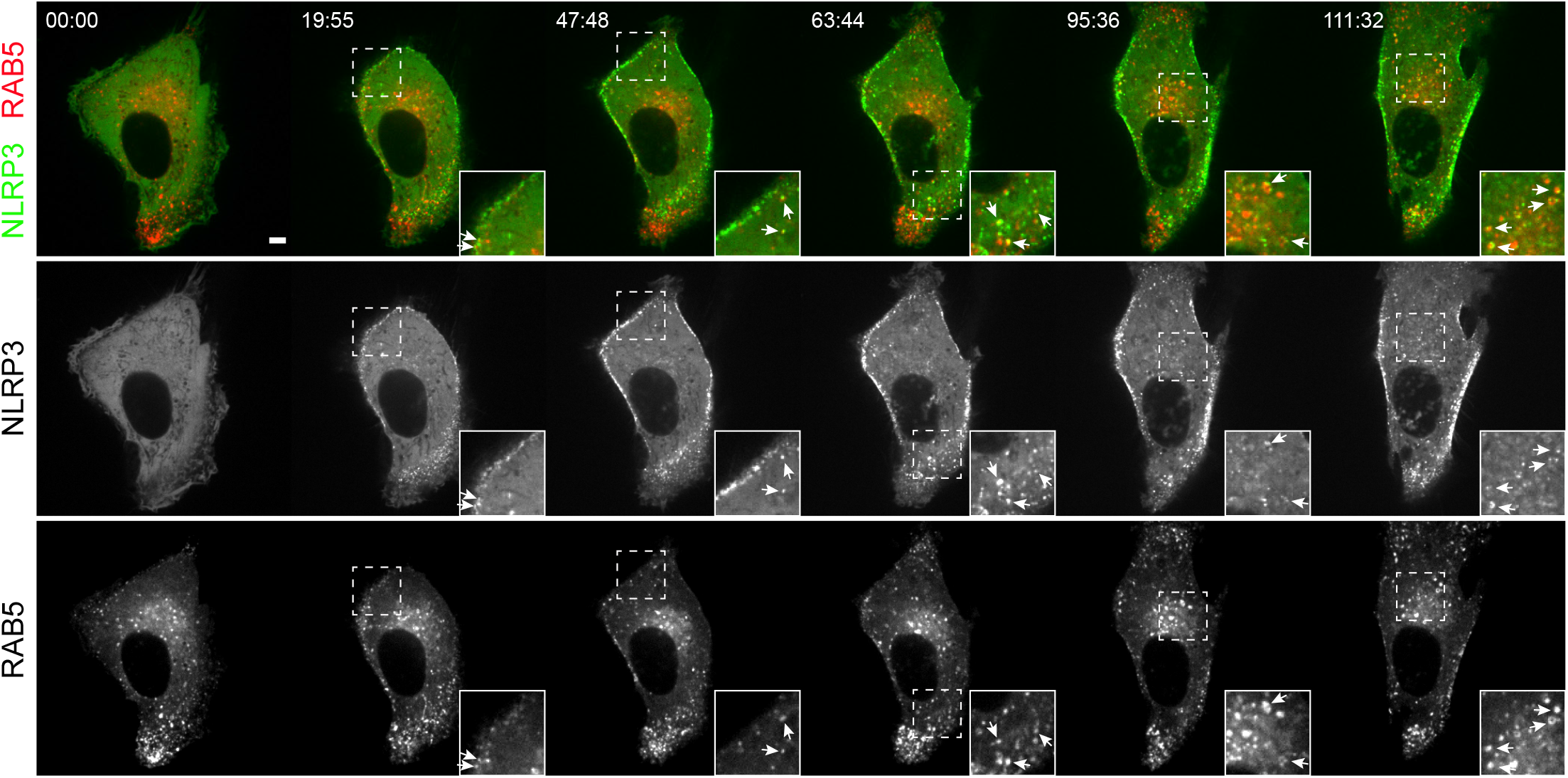
NLRP3 is recruited to vesicles of endosomal origin upon NLRP3 inflammasome activation. Spinning-disk confocal time-lapse live-video microscopy of HeLa cells stably expressing EGFP-tagged NLRP3 and mApple-tagged RAB5A. Live-video images were captured at every 239 seconds after addtion of 45 µg/ml CL097 into culture medium. Representative video stills at indicated time points are shown. Arrows indicate NLRP3 and Rab5 double-positive vesicles. Magnifications of areas are indicated with dashed squares and shown in the lower right corner of images. Arrows indicate NLRP3 and Rab5 double-positive vesicles. Scale bar: 10 µm. Images shown are representative of four independent experiments.

**Extended Data Fig.7.**
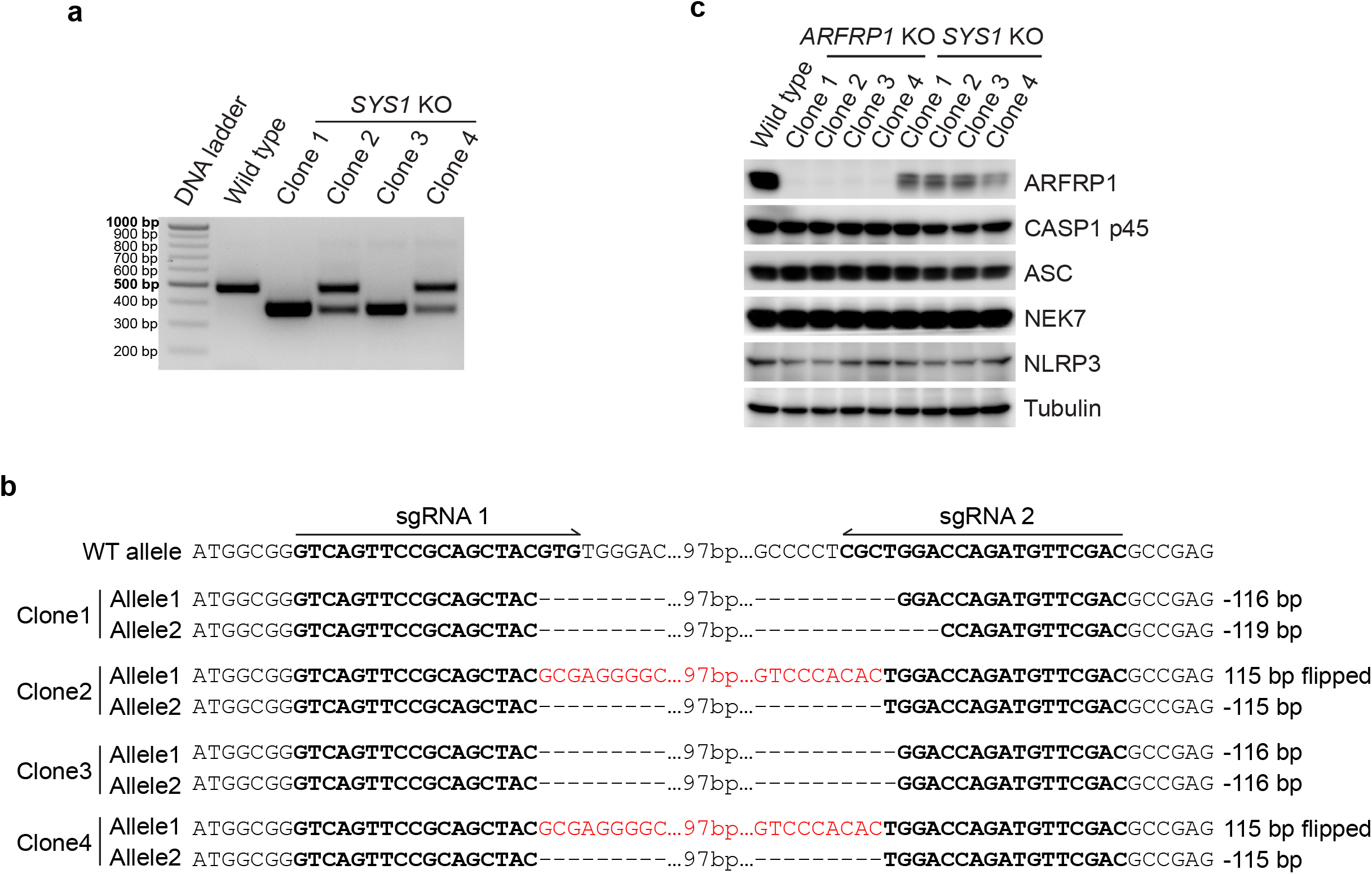
CRISPR/Cas9-mediated gene-editing allowed for generation of *ARFRP1* KO and *SYS1* KO THP-1 cells. **a**, Genotyping of *SYS1* KO THP-1 cell clones by PCR. **b**, Validation of THP-1 *SYS1* KO clones by sanger sequencing. **c**, Immunoblotting of cell lysates from Wild type, ARFPR1 KO and SYS1 KO THP-1 cells. Antibodies against ARFRP1, CASP1, ASC, NEK7 and NLRP3 were used. A antibody against Tubulin was used as a loading control. **a**,**c**, Data shown are representative of three independent experiments. **b**, sequence information was obtained from one experiment.

**Extended Data Fig.8.**
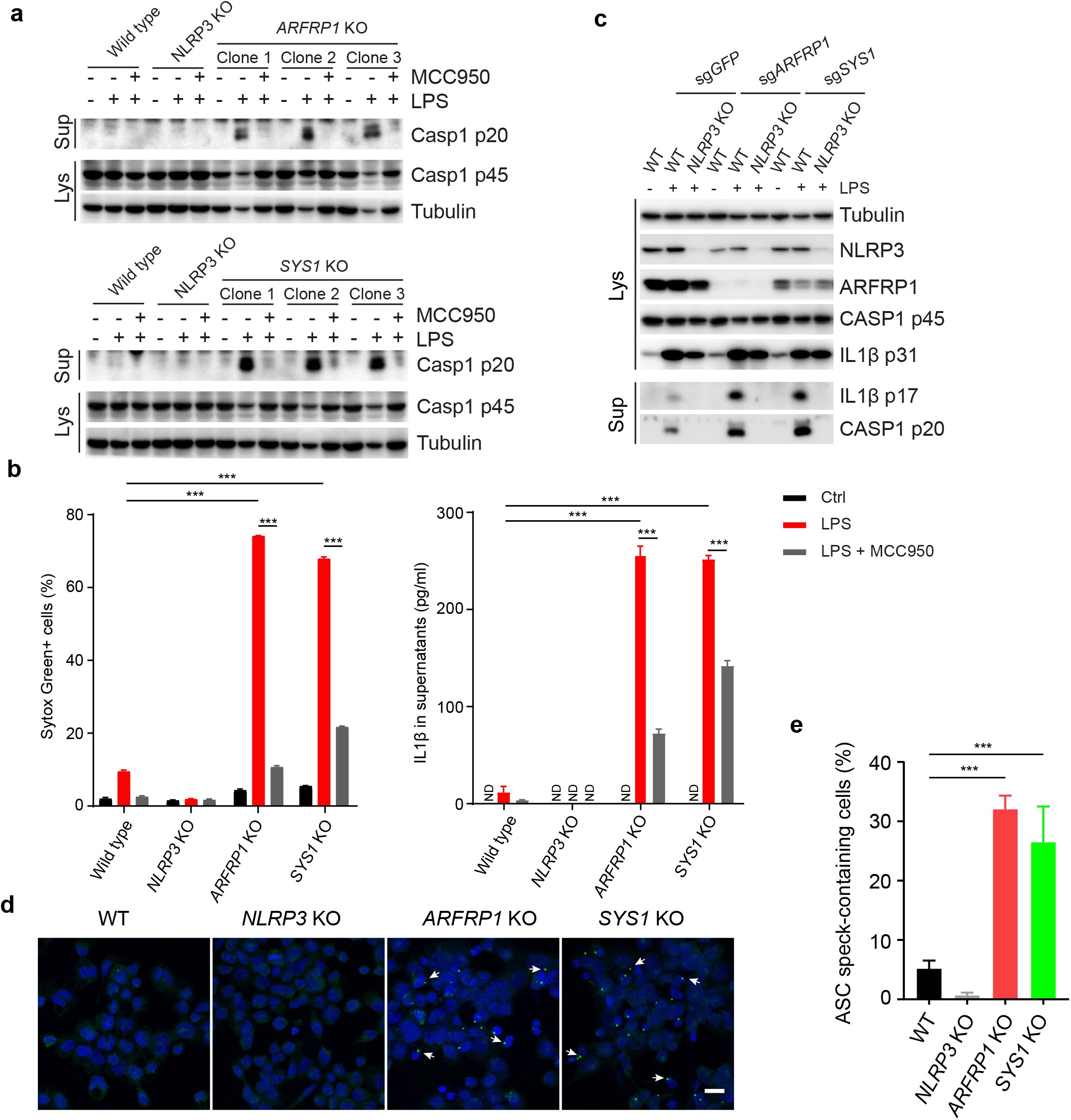
Genetic disruption of endosome-TGN retrograde transport induces activation of the NLRP3 inflammasome in response to priming only. **a**, Immunoblotting of culture supernatants and cell lysates from THP-1 cells. Wild type, *NLRP3* KO, *ARFRP1* KO (upper panel) and *SYS1* KO (lower panel) THP-1 cells were treated with 1 µg/ml LPS in absence or presence of the NLRP3 inhibitor, MCC950 (10 µM) for 2 hours. An antibody recognizing both p45 and p20 fragments of Caspase 1 (CASP1) was used. An antibody against Tubulin was used as a loading control. **b**, FACS analysis of cellular uptake of Sytox Green (left panel) and ELISA of IL1β in culture supernatants (right panel) of THP-1 cells. Wild type, *NLRP3* KO, *ARFRP1* KO and *SYS1* KO THP-1 cells were treated with 1 µg/ml LPS in absence or presence of the NLRP3 inhibitor, MCC950 (10 µM) for 2 hours. not detected (ND); *** *p* < 0.001. **c**, Immunoblotting of culture supernatants and cell lysates from THP-1 cells. PMA-differentiated wild type and *NLRP3* KO THP-1 cells depleted of ARFRP1 or SYS1 using sgRNAs were treated with 1 µg/ml LPS for 2 hours. Antibodies recognizing both p45 and p20 fragments of CASP1 as well as against IL1β, ARFRP1 and NLRP3 were used. An antibody against Tubulin was used as a loading control. **d**, Confocal microscopy of PMA-differentiated THP-1 cells. Cells were treated with 1 µg/ml LPS for 2 hours. Cells were stained with an antibody against ASC. DAPI was used to stain the nucleus. Images with z-Stacks were captured. 3-D maximal projected pictures are shown. Scale bar: 20 µm. **e**, Quantification of ASC speck-containing cells. *** *p* < 0.001. Data shown are representative of at least three independent experiments.

**Extended Data Fig.9.**
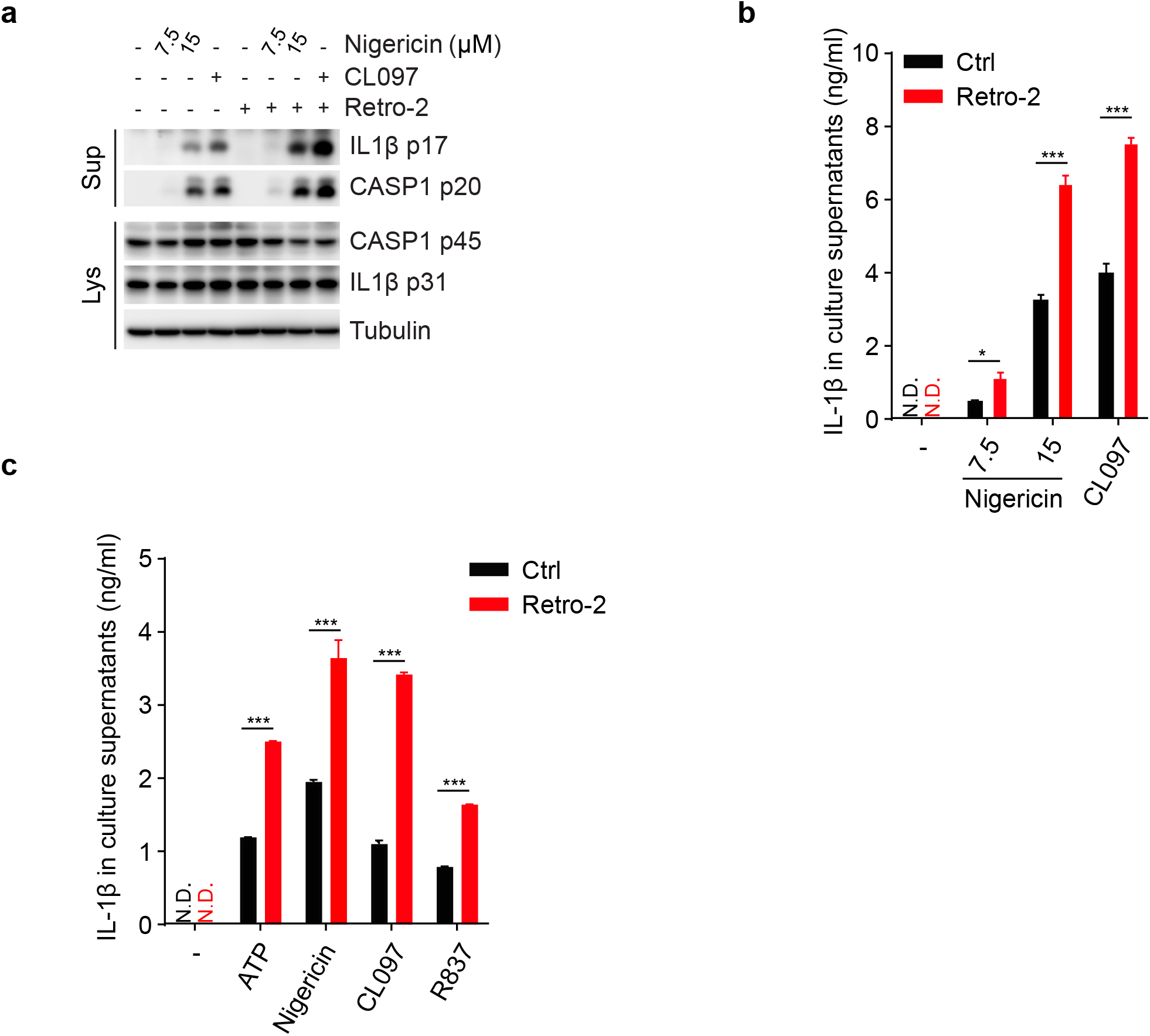
Inhibition of endosome-TGN retrograde transport by Retro-2 enhances inflammasome activation in repsonse to NLRP3 activators. **a**, Immunobloting of culture supernatants and cell lysates from PMA-differentiated THP-1 cells. Cells were pretreated with DMSO or 25 µM Retro-2 for 30 min, followed by treatment with different doses of Nigericin as indicated or 50 µM CL097 in absence or presence of 25 µM Retro-2 for 40 min. Antibodies recognizing recognizing both p45 and p20 fragments of Caspase 1 (CASP1) and agaist IL1β were used. An antibody against Tubulin was used as a loading control. **b**, ELISA of IL1β in culture supernatants from THP-1 cells. **c**, ELISA of IL1β in culture supernatants from LPS-primed BMDMs. After LPS priming, cells were pretreated with DMSO or 25 µM Retro-2 for 30 min, followed by treatment with 3 mM ATP, 7.5 µM Nigericin, 50 µM CL097 or 100 µM R837 for 30 min. not detected (ND); *** *p < 0*.*001*. Data shown are representative of three independent experiments.

**Extended Data Fig.10.**
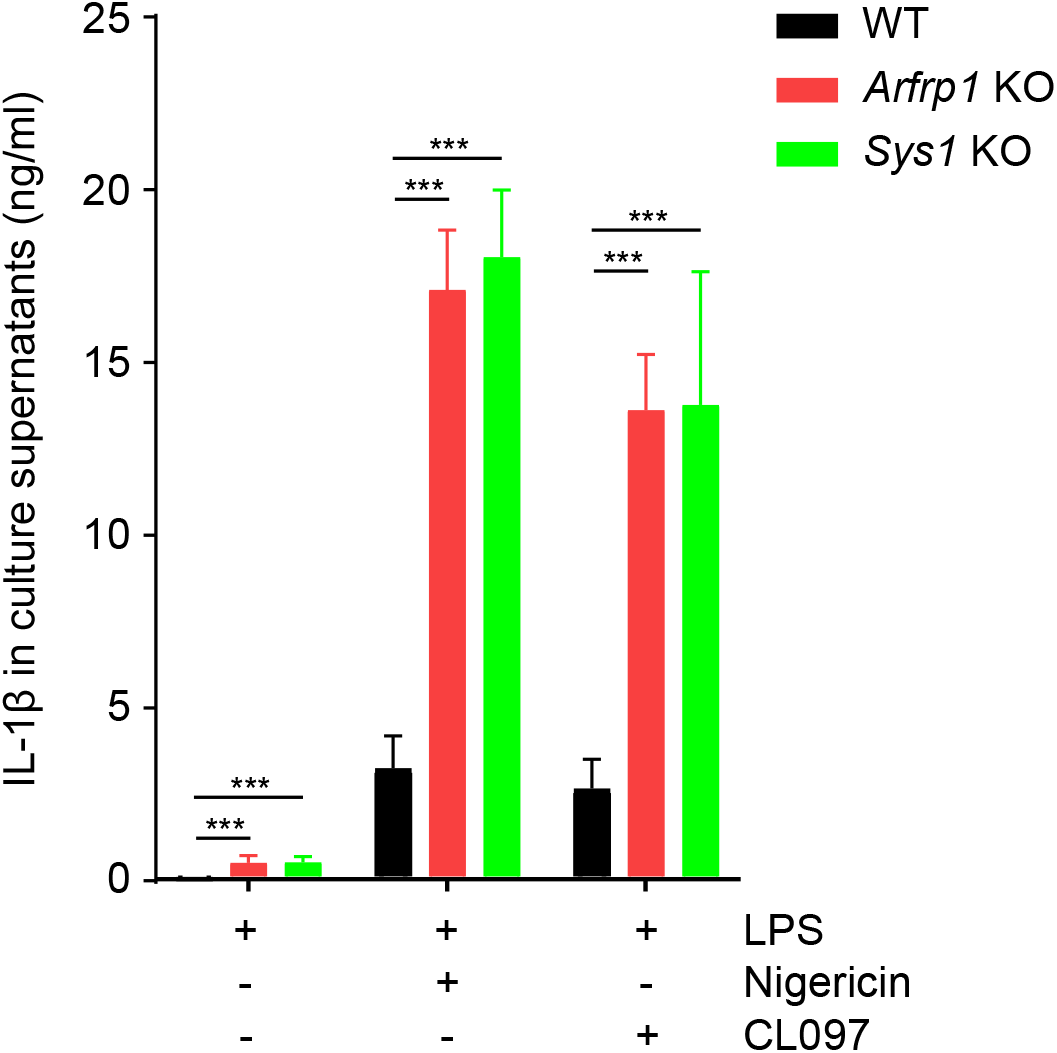
Genetic disruption of endsome-TGN retrograde transport enhances NLRP3 inflammasome activation in iBMDMS. ELISA of IL1β in culture supernatants from LPS-primed Wild type (WT), *Arfrp1* KO and *Sys1* KO iBMDMs. After LPS priming for 4 hours, cells were treated with 7.5 µM Nigericin or 50 µM CL097 for 30 min. *** *p < 0*.*001*. Data shown are representative of four independent experiments.

**Extended Data Fig.11.**
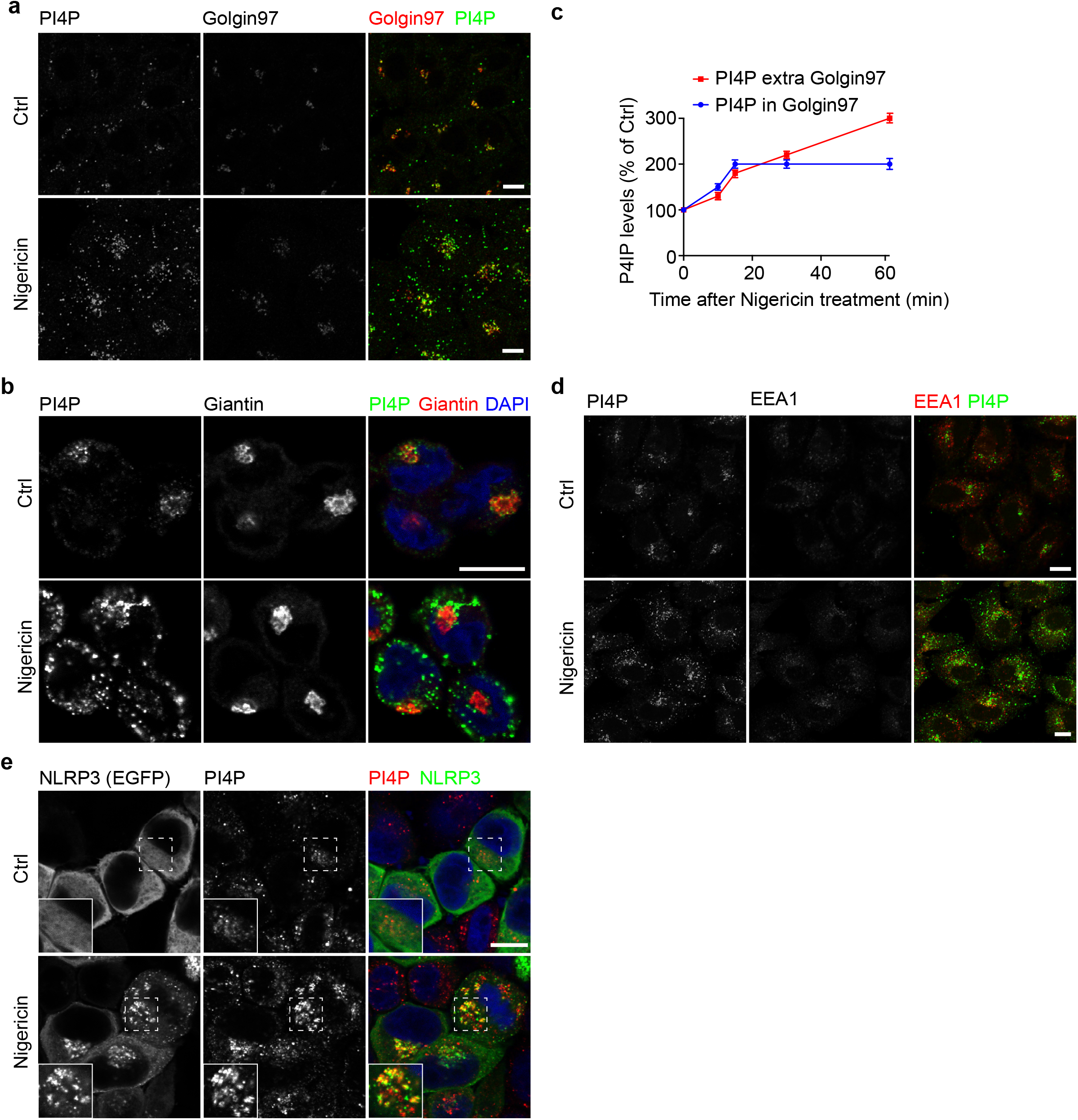
Nigericin triggers endosomal PI4P accumulation and NLRP3 co-localization with PI4P. **a**, Confocal microscopy of HeLa cells treated with vehicle or 15 μM nigericin for 60 min. After fixation, cells were stained with antibodies against PI4P and Golgin97. Scale bar: 10 μm. **b**, Quantification of PI4P levels in HeLa cells treated with vehicle or 15 μM nigericin for 0 min, 10 min, 15 min, 30 min or 60 min. Fluorescence intensity of Golgin97-colocalizing-(PI4P in Golgin97) and Golgin97-non-colocalizing (PI4P extra Golgin97) PI4P signals were quantified (100 cells per condition counted). **c**, Confocal microscopy of NLRP3 KO THP-1 cells treated with vehicle or 15 μM nigericin for 40 min. After fixation, cells were stained with antibodies against PI4P and Giantin. DAPI was used to stain the nucleus. Scale bar: 10 μm. **d**, Confocal microscopy of HeLa cells treated with vehicle or 15 μM nigericin for 60 min. After fixation, cells were stained with antibodies against PI4P and EEA1. Scale bar: 10 μm. **e**, Confocal microscopy of HeLa cells stably-expressing EGFP-tagged NLRP3. Cells were treated with vehicle or 15 μ M nigericin for 30 min. After fixation, cells were stained with an antibody against PI4P. DAPI was used to stain the nucleus. Magnifications of areas are indicated with dashed squares and shown in the lower left corner of images. Scale bar: 10 μm. **a**,**b**,**d**,**e**, Images shown are representative of at least three independent experiments.

**Extended Data Fig.12.**
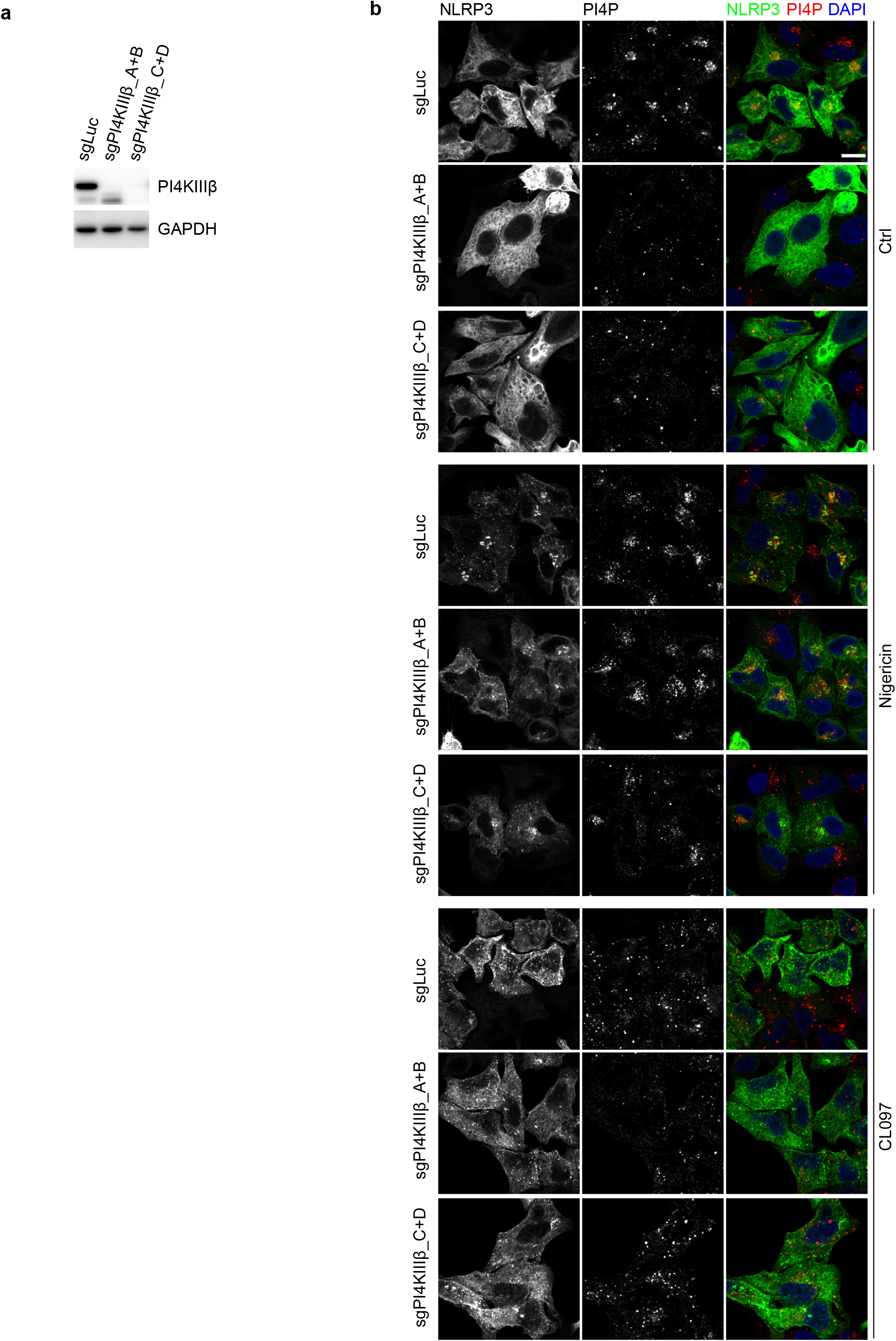
Deletion of PI4KIIIβ does not affect PI4P accumulation in and NLRP3 recruitment to endosomes. **a**, Immunoblotting of cell lysates from EGFP-tagged NLRP3-stably expressing HeLa cells co-expressing sgRNAs targeting Luciferase (sgLuc) or PI4KIIIβ (sgPI4KIIIβ). Two different pairs of sgRNAs were used to delete PI4KIIIβ (pairs A+B and C+D as indicated in the methodology). An antibody against PI4KIIIβ was used. An antibody against GAPDH was used as a loading control. **b**, Confocal microscopy of HeLa cells stably expressing EGFP-tagged NLRP3 co-expressing sgRNAs targeting Luciferase (sgLuc) or PI4KIIIβ (sgPI4KIIIβ). Cells were treated with vehicle, 10 μM Nigericin for 30 min or 45 μg/ml CL097 for 80 min and then stained with an antibody against PI4P. DAPI was used to stain the nucleus. Scale bar: 10 μm. Data shown are representative of three independent experiments.

**Extended Data Fig.13.**
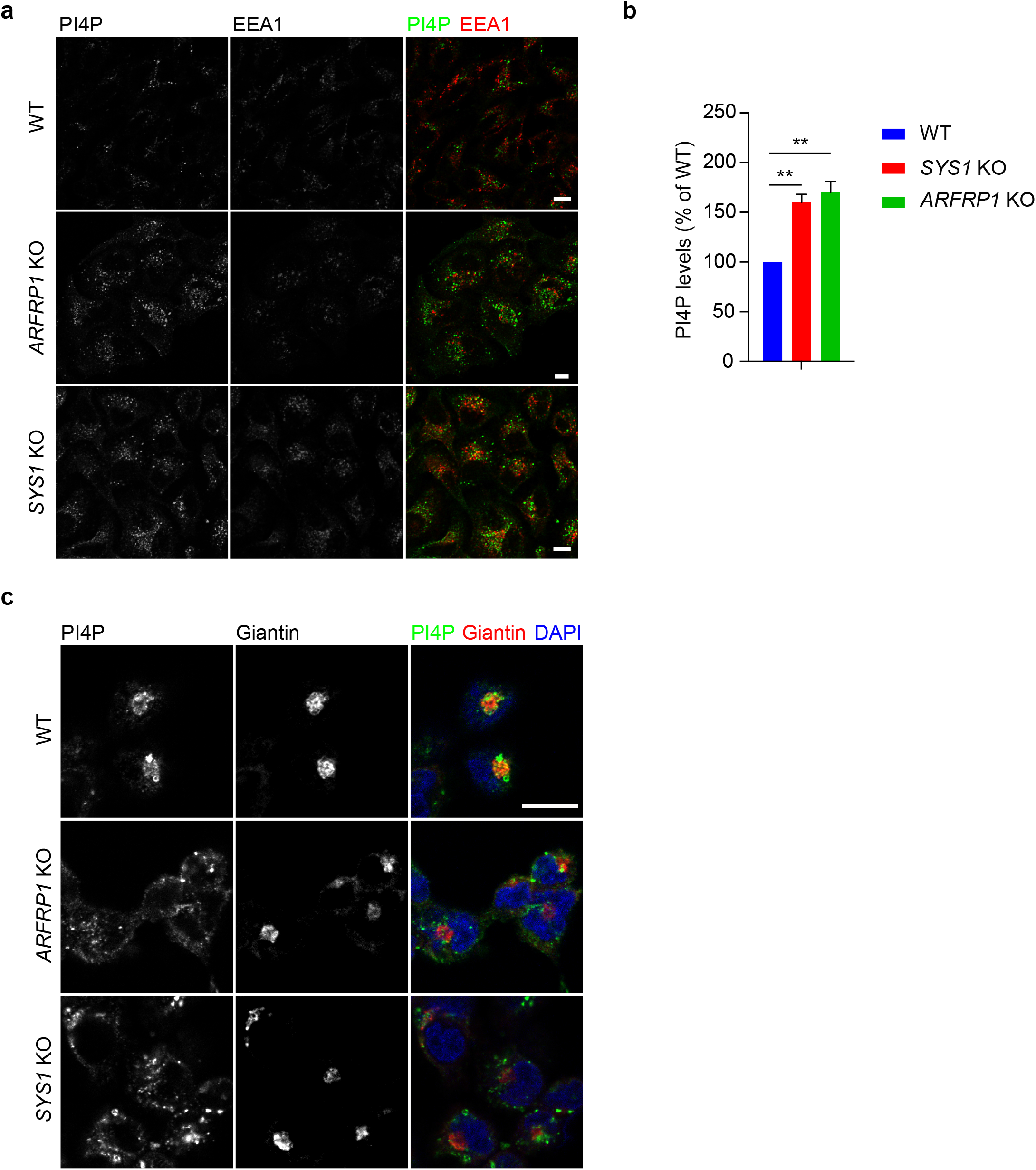
Disruption of endsome-TGN retrograde transport leads to PI4P accumulation on endosomes. **a**, Confocal microscopy of Wild type (WT), *ARFRP1* KO and *SYS1* KO HeLa cells. Cells were co-stained with antibodies against PI4P and EEA1. Scale bar: 10 μm. **b**, Quantification of PI4P levels in cells. ** *p* < 0.01 (100 cells per condition counted). **c**, Confocal microscopy of WT, *ARFRP1* KO and *SYS1* KO THP-1 cells. Cells were co-stained with antibodies against PI4P and Giantin. DAPI was used to stain the nucleus. Scale bar: 10 μm. **a**,**c**, Images shown are representative of three independent experiments.

## Methods

### Methods Mice

Mice with targeted alleles for *Arfrp1* (*Arfrp1*^*fl/fl*^) were described previously (*25*) and were provided by Dr. Annette Schürmann (German Institute of Human Nutrition Potsdam-Rehbruecke, Germany). *Arfrp1*^*fl/fl*^ mice on C57BL/6J background were crossed with *LysM-Cre* mice to obtain myeloid-specific *Arfrp1* knockout mice. Mice were housed under specific pathogen-free conditions with controlled temperature on a 12-h light/dark cycle with unrestricted access to water and standard laboratory chow. Maintenance and animal experimentation were in accordance with the local ethical committee (Com’Eth) in compliance with the European legislation on care and use of laboratory animals (La cellule AFiS (Animaux utilisés à des Fins Scientifiques): APAFIS#30865-2021040115389039 v3). No exclusion of animals used for experiments was performed. Healthy mouse littermates were chosen randomly according to their genotypes.

### Reagents

Nigericin sodium salt (catalogue no. N7143), Adenosine 5’-triphosphate disodium salt (ATP) (A2383), Retro-2 (SML1085), STxB (SML0562), Phorbol 12-myristate 13-acetate (P8139) and Lipopolysaccharides from *Escherichia coli 055:B5* (L2880) were purchased from Sigma-Aldrich. Pam3CSK4 (tlrl-pms), Imiquimod (tlrl-imq) and CL097 (tlrl-c97) were obtained from InvivoGen. SYTOX™ Green Nucleic Acid Stain (S7020) was purchased from Thermo Fisher Scientific. Cy3-STxB was generated by conjugating Cy3 dye onto STxB according to the manufacture manual of CY®3 Ab Kit (PA330000, Sigma-Aldrich).

### Plasmids

pX330-U6-Chimeric_BB-CBh-hSpCas9 (Plasmid #42230) and lentiCRISPR v2 (Plasmid #52961) were purchased from Addgene. EGFP-tagged mouse *NLRP3* was cloned into pBOB empty vector using ligation-independent cloning (LIC) as described previously (*26*). pEGFP-N1 hSYS1 and pEGFP-N1 hARFRP1 (WT, Q79L, T31N) were described previously (*17*). Flag-tagged hSYS1, EGFP-tagged hARFRP1 (WT, Q79L, T31N) sequences were amplified by PCR and were cloned into pBOB empty vector. The pX330-P2A-EGFP/RFP plasmids were generated by inserting P2A-EGFP/RFP sequence into EcoR I-digested pX330-U6-Chimeric_BB-CBh-hSpCas9 before stop codon using LIC. Small guide RNA (sgRNA) sequences targeting human *ARFRP1* gene or human *SYS1* gene were inserted into Bbs-I-digested pX330-P2A-EGFP/RFP plasmids through ligation by T4 DNA ligase. SgRNA sequences targeting mouse Arfrp1 gene or mouse SYS1 gene were inserted into Bbs-I-digested lentiCRISPR v2 through ligation by T4 DNA ligase.

### Cell culture

All mammalian cells were cultured at 37 °C, 5% CO2. Cell lines used in this study are not listed in these databases of commonly misidentified cell lines maintained by ICLAC. THP-1 cells (ATCC) were grown in RPMI 1640 containing 10% fetal bovine serum, 10 mM HEPES, 2.5 g/l glucose, 1 mM sodium pyruvate and gentamycin. Hela cells were grown in DMEM (4.5 g/l glucose) containing 10% fetal bovine serum, penicillin and streptomycin. HEK293t cells (DKFZ Heidelberg) were grown in DMEM containing 1 g/ml glucose, 10% fetal bovine serum, penicillin and streptomycin. The generation of THP-1 NLRP3 knockout cells were described previously (*26*). mApple-tagged RAB5 stably-expressing HeLa cell line was provided by Dr. Anne Spang (BIOZENTRUM, University of Basel, Switzerland). Immortalized bone marrow-derived macrophages (iBMDMs) cell line was obtained from Dr. Eicke Latz (University of Bonn, Germany) and was grown in DMEM (4.5 g/l glucose) containing 10% heat-inactivated fetal bovine serum, penicillin and streptomycin. Primary BMDMs were obtained by differentiating bone marrow progenitors from the tibia and femur in RPMI 1640 containing 50 ng/ml recombinant hM-CSF (11343113, Immunotools), 10% heat-inactivated fetal bovine serum, penicillin and streptomycin for 7 days. BMDMs and peritoneal macrophages were seeded 1 day before experiments using RPMI 1640 containing 10% heat-inactivated fetal bovine serum, penicillin and streptomycin. Cell cultures were negative for mycoplasma contamination. THP-1 cells and HEK293t cells have been authenticated using Short Tandem Repeat (STR) performed by LGC Standards, UK.

For treatment of cells, BMDMs were primed with 1 µg/ml LPS for 4 h, followed by the treatment of ATP, Nigericin, Imiquimod and CL097; while THP-1 cells were differentiated by PMA (100 nM) treatment for 3 h followed by overnight incubation with fresh medium before experiments. For Retro-2 treatment, the cells were pretreated with 25 µM Retro-2 for 30 min before the treatment of cognate stimuli in present of Retro-2.

For the packaging of lentivirus, 12 µg of Lenti-mix (3 µg pVSVG, 3 µg pMDL and 3 µg pREV) plus 12 µg of gene of interest expressing plasmid were transfected into HEK293t cells (10-cm plate) using Lipofectamine 2000. After 48 h, the supernatants were collected and filtered using 0.45 µm Millex-HV Syringe Filters and kept at -80 °C. BMDMs and THP-1 cells were infected in fresh medium containing 1 µg/ml polybrene (sc-134220, Santa Cruz) and 25% lentivirus-contained supernatant.

### Gene disruption using CRISPR/Cas9 genome editing system

To generate the *ARFRP1* KO and *SYS1* KO THP-1 cell lines, two small guide RNA sequences for each gene were designed using Website https://www.benchling.com/ and cloned into pX330-P2A-EGFP and pX330-P2A-RFP, separately, through ligation using T4 ligase. 1.0 × 10^6^ THP-1 cells were transfected with mixture of two sgRNAs-expressing plasmids (0.5 µg each) using X-tremeGENE™ 9 DNA Transfection Reagent (6365779001, Roche) according to manufacturer’s manual. 24 h after transfection, GFP and RFP double-positive cells were enriched by FACS sorting (BD FACS Aria II). Single cell colonies were obtained by seeding them into 96-well plates by series of dilution. Obtained *ARFRP1* KO and *SYS1* KO single-cell clones were validated by immunoblotting or Sanger sequencing of PCR-amplified targeted fragment after cloned into pUC57 vector. The following primers were used for PCR amplification of SYS1 targeted fragment: 5’-CGAATGCATCTAGATATCGGATCCACTCTGAGAATGGGTCTGTCTGCCC-3’ and 5’-GCCTCTGCAGTCGACGGGCCCGGGTTCCCAGGGCTACAAAGAAAGAAGG-3’.

To generate *Arfrp1* KO and *Sys1* KO iBMDM cell lines, two sgRNAs for each gene were designed and cloned into lentiCRISPR v2. Lentiviruses were produced as described above. After 24 hours after lentiviral infection, sgRNAs expressing-iBMDMs were selected by treatment with 2 µg/ml puromycin for 24 hours. Single cell colonies were obtained by seeding them into 96-well plates by series of dilution. Obtained Arfrp1 knockout and Sys1 knockout single-cell clones were validated by immunoblotting or Sanger sequencing.

To generate EGFP-tagged NLRP3 stably-expressing HeLa cells depleted of PI4KIIIβ, two sets of sgRNAs (two sgRNAs in each set) were designed and cloned into lentiCRISPR v2. Lentiviruses were produced as described above. EGFP-tagged NLRP3 stably-expressing HeLa cells were infected with sgRNA-expressing lentivirus as indicated. After 24 hours after lentiviral infection, sgRNAs expressing-cells were selected by treatment with 2 µg/ml puromycin for 24 hours. After two splitting, cells were used for experiments.

The sequences of sgRNAs used in this study are listed below:

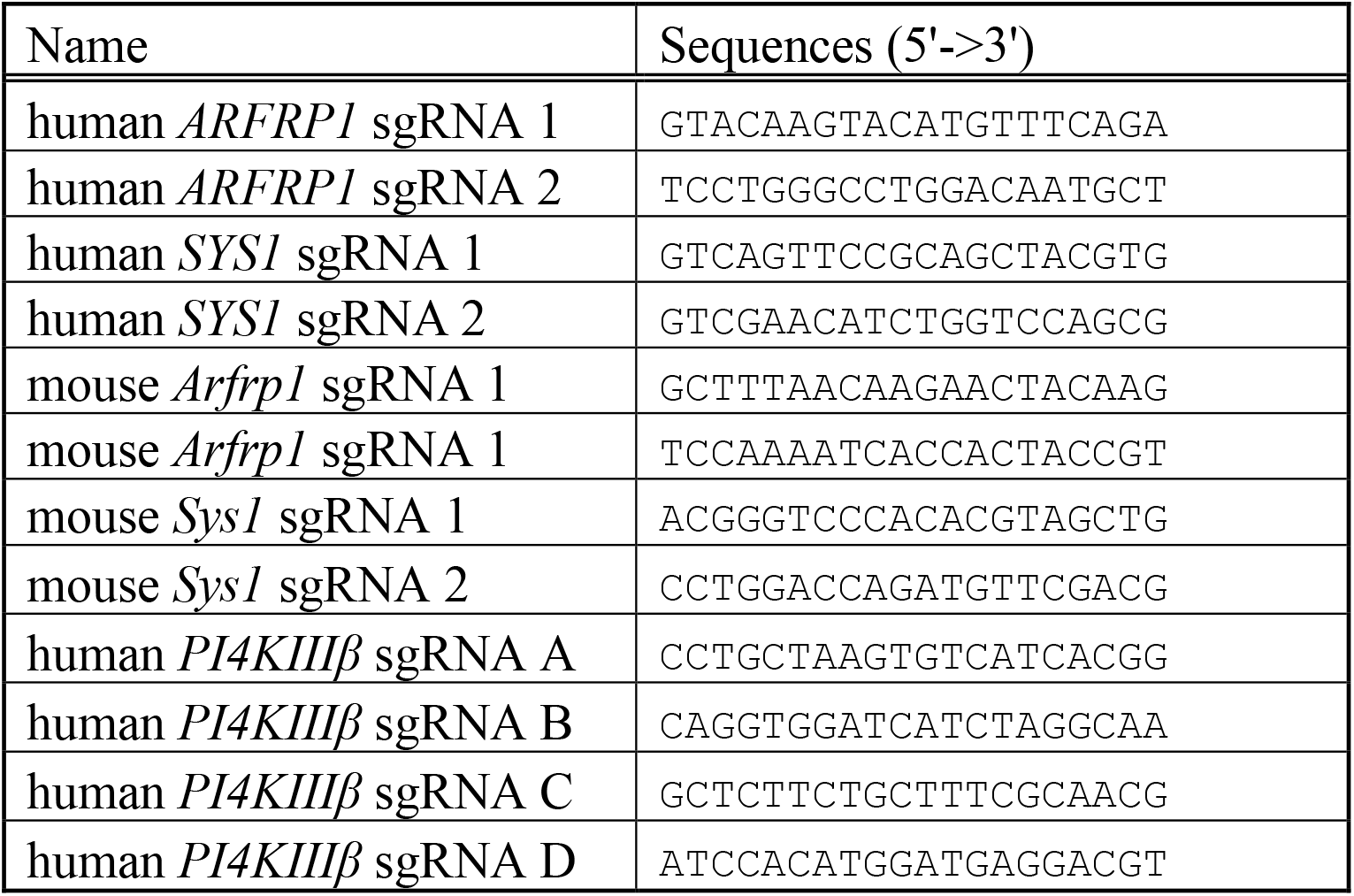

### Immunoblotting

After treatments, cell culture supernatants and cell lysates were collected for immunoblotting analysis. Cell lysates were collected in 1× SDS sample buffer or in 1×RIPA buffer (50 mM Tris-HCl (pH 7.5), 150 mM NaCl, 1% Triton X-100, 1 mM EDTA, 1 mM EGTA, 2 mM Sodium pyrophosphate, 1 mM NaVO4 and 1 mM NaF) supplemented with protease inhibitors cocktail. The immunoblot were prepared using Tris-Glycine SDS-PAGE. For the supernatants, the proteins were extracted using methanol-chloroform precipitation, separated by Tricine SDS-PAGE and analyzed by immunoblotting. The immunoblots were probed overnight at 4 °C or 1 hour at room temperature with anti-human IL-1β antibody (AF-201-NA, R&D Systems); anti-human Caspase-1 antibody (06-503, Merck Millipore); anti-mouse IL-1β antibody (5129-100, BioVision); anti-mouse Caspase-1 p20 (AG-20B-0042-C100, AdipoGen); anti-ASC antibody (sc-22514, Santa Cruz Biotechnology), anti-NLRP3 antibody (G-20B-0014-C100, AdipoGen); anti-PI4KIIIβ antibody (611816, BD Bioscience); anti-GAPDH (G9545, Sigma-Aldrich); anti-Tubulin (T9026, Sigma-Aldrich); anti-ARFRP1 (PA5-50606, ThermoFisher Scientific) and anti-Flag antibody (F1804, Sigma-Aldrich).

### Immunofluorescence microscopy

After treatments, cells plated on coverslips (9-15 mm) were fixed in 4% paraformaldehyde for 15 min at room temperature. Cells were permeabilized in phosphate-buffered saline (PBS) containing 0.1% Saponin for 10 min. After blocking with 10% normal goat serum in PBS for 1 h, cells were incubated with primary antibodies for 1 h at room temperature. After incubation with secondary antibodies for 1 h at room temperature, cells were stained with DAPI and mounted with ProLong™ Gold Antifade Mountant (P36934, ThermoFisher Scientific). Intracellular PI4P staining was performed as previously described in Hammond et al., 2009 (*27*). Images were acquired below pixel saturation using Confocal Laser Scanning Microscope Leica TCS SP8 (Leica microsystem).

Antibodies used in this study for immunofluorescence staining are listed below:

| Name of antibody | Provider | Catlog No. | Dilution |
| --- | --- | --- | --- |
| Rabbit anti-TGN46 | ProteinTech | 13573-1-AP | 1/300 |
| Sheep anti-Human TGN46 | Bio-rad | AHP500GT | 1/50 |
| Rabbit anti-GCC2 antibody | Sigma-Aldrich | HPA035849 | 1/200 |
| Rabbit anti-GCC1 antibody | Sigma-Aldrich | HPA021323 | 1/100 |
| Mouse anti-human p230 | BD Bioscience | 611280 | 1/100 |
| Mouse anti-Golgin97 | ThermoFisher Scientific | A-21270 | 1/100 |
| Mouse anti-EEA1 | BD Bioscience | 610456 | 1/100 |
| Rabbit anti-EEA1 | Cell Signaling Technology | 3288S | 1/100 |
| Mouse anti-PI4P | Echelon Biosciences | Z-P004 | 1/100 |
| Rabbit anti-Golgin97 | Home-made | Venditti *et al.*, 2019 (28) | 1/100 |

### Live-video imaging

Hela cells stably-expressing EGFP-tagged NLRP3 and mApple-tagged RAB5A were seeded into Nunc™ Lab-Tek™ 8-well Chambered Coverglass (155411, ThermoFisher Scientific) on the day before experiments. On the second day, cells (with 50-60% confluence) were treated with 10 μM nigericin or 45 μg/ml CL097. Live images with *z*-stacks were acquired at every 3 mins using Nikon Inverted confocal spinning-disk microscope.

### Uptake and trafficking of CI-M6PR antibody and Cy3-labelled StxB

CI-M6PR antibody uptake was performed as previously described (*29*). Cells were treated with 10 μM nigericin or 45 μg/ml CL097 in serum-free medium. At 5 min after nigericin addition or 20 min after CL097 addition, an anti-CI-M6PR antibody (10 μg/ml) (MA1-066, Thermol fisher Scientific) was added into culture medium. Cells were fixed at 40 min after the incubation of CI-M6PR antibody and co-stained with antibodies against GCC1 or EEA1. For StxB uptake, Cy3-conjugated StxB was added into culture medium at 5 min after nigericin (10 μM) addition or 20 min after CL097 (45 μg/ml) addition. Cells were fixed at 20 min or 40 min after the incubation of Cy3-conjugated StxB and stained with cognate antibodies. Images were acquired using Confocal Laser Scanning Microscope Leica TCS SP8 (Leica microsystem).

### Analysis of Sytox Green uptake using flow cytometry

After treatment, cells were detached with 5 mM EDTA and incubated with SYTOX™ Green (S7020, ThermoFisher Scientific) at 1/8000 dilution. Cells were then analyzed using BD FACS Celesta™ Cell Analyzer (BD Bioscience). Percentage of Sytox Green-positive cells was analyzed using FlowJo software.

### LPS-induced endotoxemia in mice

*Arfrp1*^*fl/fl*^ and *Arfrp1*^*fl/fl*^;*LysM-Cre* mice at 6-8 weeks old were used. Mice were peritoneally injected with 15 mg/kg LPS. For the analysis of serum cytokines, blood was collected at 3 hours after LPS injection. For survival curves, 2.5 mg/kg Buprenorphine was peritoneally administrated every 6 hours and the survival of mice was monitored every 6 hours.

### Measurement of cytokines using ELISA

Human IL-1β, Mouse IL-1β and TNFα and IL-6 in cell culture supernatants or serum were measured using Human IL-1 beta/IL-1F2 DuoSet ELISA (DY201, R&D Systems), Mouse IL-1 beta/IL-1F2 DuoSet ELISA (DY401, R&D Systems) or Mouse TNF-alpha DuoSet ELISA (DY410, R&D Systems), separately, according to the manufacturer’s protocols.

### Image quantification

Image processing and quantification was performed using Fiji software (National Institutes of Health) (*30*).

For the quantification of StxB and CI-MPR on endosomes and Golgi, to define ROIs, the EEA1 (for endosomes) and GCC1 (for Golgi) channels were thresholded. Defined ROIs was superimposed onto the channel of interest (StxB and CI-MPR). Fluorescence in the defined ROIs were measured by recording “RawIntDen” (the sum of the values of the pixels in the selection). The percentage of StxB and CI-MPR in defined ROIs were calculated through dividing “RawIntDen” in the defined ROIs by “RawIntDen” of the whole cell. A single mid-z stack image was analyzed per cell.

For the quantification of PI4P levels, a mask using the TGN marker Golgin-97 was generated for each cell, and the mean intensities of both PI4P and Golgin-97 were measured in those regions. After background subtraction, the PI4P values were normalized singularly using their own Golgin-97 values for each time points. For PI4P level measurement in WT, *ARFRP1* KO and *SYS1* KO HeLa cells, mean intensities of both PI4P and EEA1 were calculated individually for each cell. Then, similarly, the PI4P values were normalized using their own EEA1 values, after background subtraction.

### Statistical analyses

Preliminary experiments were performed and sample size was determined based on generally accepted rules to test preliminary conclusions reaching statistical significance, where applicable. Unless specified, statistical analyses were performed with the *t*-test using GraphPad Prism (GraphPad Software). For serum cytokine levels, statistical analyses were performed with the *Mann Whitney* test. For survival curves, statistical analysis was performed with the *Log-rank (Mantel-Cox)* test.

